# IFNα, a potential biomarker for stress vitiligo risk

**DOI:** 10.1101/151571

**Authors:** Huali Wu, Ting Wang, Minxuan Cai, Mengsi Fu, Fengfeng Ping, Ling He, Xiaohong An, Zhixiang Shi, Zhenjiang Xia, Jing Shang

## Abstract

Neural hypothesis has become an important aspect of vitiligo, yet without corresponding diagnostic indicators. We preliminarily found 32 cases of vitiligo patients with certain aggregation of mental factors. In peripheral blood mononuclear cells (PBMCs) of these patients, transcriptome analyses revealed that the circulation expression of a type I interferon (IFN-I)-dependent genes was induced. Also, serum IFNα was elevated in vitiligo patients with depression. Therefore, our hypothesis is whether IFNα levels predict the occurrence of psychiatric vitiligo. Through the establishment of stress-induced depigmentation model, serum IFNα also showed increase. Intracerebroventricular and subcutaneous IFNα injection can both elicit not only depressive behavior but also vitiligo-like characteristics. Mechanistically, central IFNα induces the release of dorsal root ganglion (DRG) substance P (SP) to inhibit melanogenesis. Peripheral IFNα disturbs cutaneous-neuro-endocrine microenvironment. Type I IFN (IFNα) pathway-related genes in stress vitiligo were significantly discriminating from non-stress vitiligo, while that of type II IFN pathway was not.

## Introduction

In the clinical study of new drugs, the research and application of biomarkers are gradually focused. Based on biomarker-based precision drug treatment, patients were screened and grouped according to the corresponding biomarkers, and the response rate of the specific patient population to the drug would be higher. Accordingly, the US FDA has introduced a new regulatory guideline: clinical research on the application of anti-tumor drug should provide biomarker diagnostic kit at the same time. These biomarkers could then be applied to the general population to ensure patients receive appropriate medication. Vitiligo, a puzzling disease with complex pathogenesis, is not only a local pigment disappearance but a systemic disease. There are several theories, involving neural, autoimmune, and biochemical mechanisms. Because patients show the same signs and symptoms, clinical therapeutic strategies are not very well distinguished. Currently, dermatologists generally classify patients into autoimmune diseases (1). The extensive use of immunomodulators has certain effects, which are not significant. Therefore, there may exist other therapeutic approaches targeting different pathological mechanisms such as neural. In recent years, neural hypothesis is most widespread and has become an important aspect of vitiligo incentives. However, in clinical there are no corresponding diagnostic indicators and treatment drugs.

At present, due to the lack of rapid and effective diagnostic indicators used to accurately classify vitiligo patients, clinical treatments often blindly choose surgery or immunosuppression by combination of light therapy. China Vitiligo Treatment Consensus (2014 Edition) categories include: (1) VIDA score, (2) vitiligo Kobner test (KP), (3) clinical characteristics, (4) Woods lamp observation, (5) Skin CT supplementary diagnosis, and (6) skin pathology. The above four points can comprehensively assess the progression of the disease. 2011 Vitiligo Global Symposium (VGICC) revealed: lack of judgment indicators. In recent years, vitiligo progression biomarkers have been given more and more attention and a lot of useful exploration has been carried out. For example, in 2017, Reinhart et al reported that S100B could be a biomarker that indicates the progression of vitiligo (2). In 2016, Xiang LH et al found that serum CXCL10 may be a new molecular marker that detect disease progression and guide the treatment (3). However, from the aspect of vitiligo etiology, molecular markers, used to classify patients, have not been studied.

The psychogenic theory receives more and more attention. Clinical observations have shown that approximately one-third of patients may have psychiatric comorbidity and the prevalence of depression in vitiligo patients was 39% in a QoL study (4). A series of experimental studies have also shown that mental stress can cause mouse pigment loss (5-7). Therefore, it is undeniable that vitiligo is a typical physical and mental illness, which is closely related with mental and neurological factors. With the rapid development of biomedicine, new biomarkers are continually found for neurological diseases. For example, the levels of tau total protein, phosphorylated tau protein and beta amyloid Aβ42 in cerebrospinal fluid can be used as biomarkers for the diagnosis of early Alzheimer’s disease (8). Cerebrospinal fluid or blood levels of IFNα in neural lupus erythematosus (SLE) patients were significantly increased (9), suggesting that IFNα level can predict the occurrence of neurological SLE. In vitiligo, the IFN factors are abnormally changed, with significant increase of IFNγ and extremely low expression of IFNα in skin lesion (10). IFNγ in skin can regulate the maturity of melanocyte, stimulate the secretion of IL6 and IL8 in keratinocytes and finally inhibit the melanogenesis (11). It can be concluded that IFNγ can directly participate in the vitiligo onset through a local action. Literature survey has pointed out that IFNα may activate the downstream signaling network, and modulate the progression of vitiligo (12). Clinically, successive administration of IFNα is able to induce vitiligo phenotype and depression (13-15). The depression rate is as high as 30~45% (14) and the treatments have to be interrupted. IFNα in the serum of vitiligo patients is significantly increased, yet it is extremely low in skin. Under stress conditions, our transcriptome data analyses showed that the circulation expression of a type I interferon (IFN-I)-dependent genes of vitiligo patients was largely induced (**Figure 1**). However, expression of IFN-II-dependent genes slightly played the effect (**Figure 1**). Here, we suppose that the melanogenic effects induced by IFNα are systematic and closely connected with the nervous system. Some researches suggest that IFNα can induce brain dysfunction directly or indirectly, leading to anxiety, depression and other neuropsychiatric diseases (16-18). Only a little IFNα can pass BBB and it can not cause depression when IFNAR is blocked in central and peripheric regions (18). Therefore, both central and peripheric IFNα can induce neuropsychiatric diseases via IFNAR. The endogenic IFNα in brain can lead to neuroimmunoreactive dysfunction, and then activate the endocrine and immunity system, causing several systematic diseases, such as systemic lupus erythematosus(SLE) (9). Clinical studies suggest that IFNα level in the brains of neural SLE patients is significantly higher than that in blood. The IFNα level in the blood of vitiligo patients is up-regulated and administration of IFNα can induce vitiligo phenotype and depression. IFNα mainly mediate depression via nervous system, hypothalamic–pituitary–adrenocortical (HPA) axis and immune system. Vitiligo is not just local depigmentation but a systematic disease. From all the above, we suppose that IFNα may be a biomarker of psychogenic vitiligo (induced by neural factors). IFNα level in blood can divide the patients into psychogenic and non-psychogenic vitiligo groups, which would provide significant evidence for subsequent treatment and greatly improve the cure rate.

**Figure 1.**
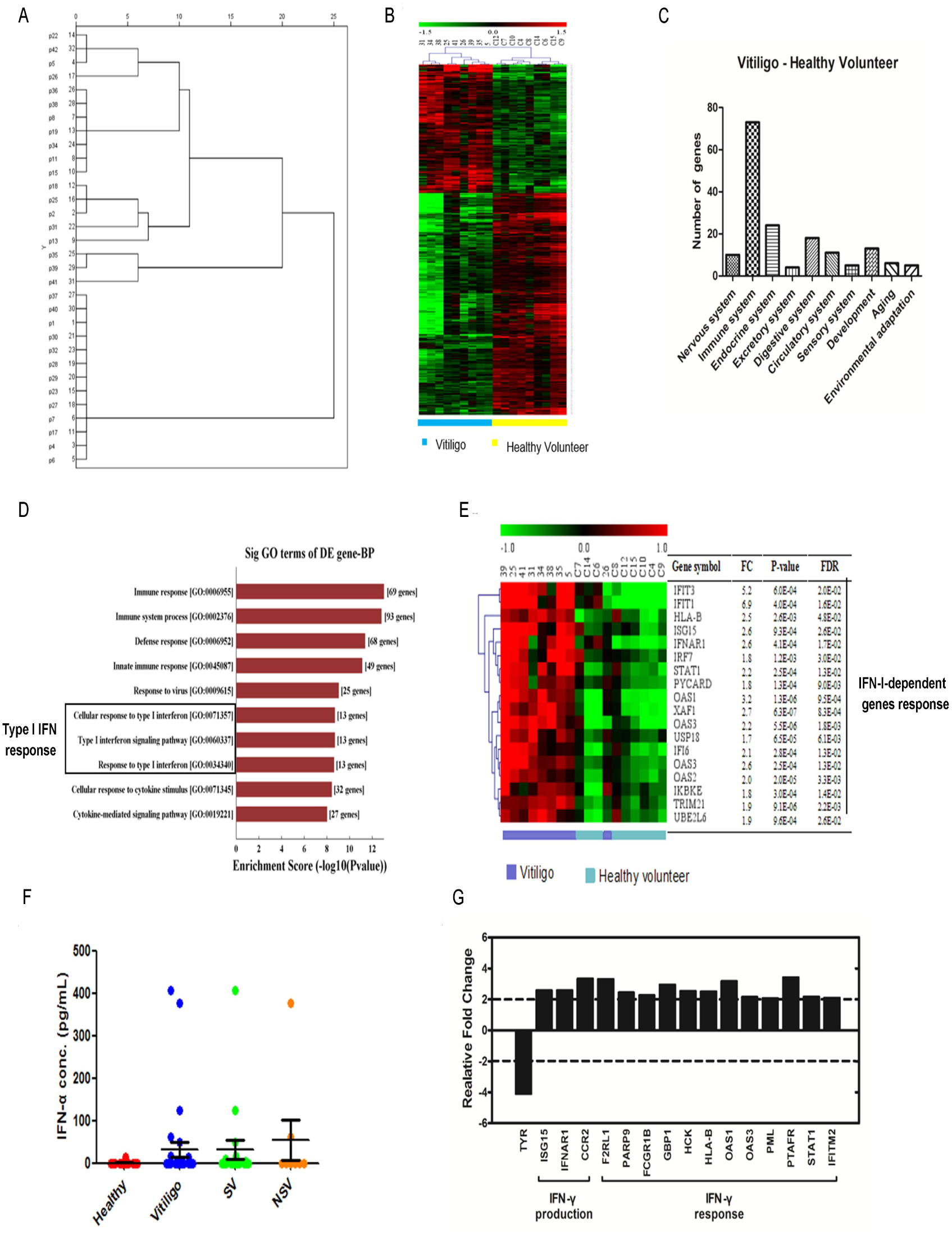
Vitiligo patients have a certain aggregation of mental factors and a distinct IFN-I-dependent signal expression profile in the blood. (A) By combining the HAMD and the HAMA 14 scales, we clustered a total of 18 people with mental disorders of vitiligo. (B) Heat map, vitiligo patients have a distinct transcriptome profile in the blood. The clustering correctly reflects the experimental design and the differential expression pattern between the study groups. (C) Bar graphs of differentially expressed genes in nervous system, immune system, endocrine system, excretory system, circulatory system, etc. (D) Most significantly enriched groups for the differentially regulated mRNAs relating to biological processes (BP). (E) Heatmap and cluster dendrogram of IFN-I dependent genes. (F) The serum IFN-α level of vitiligo patients. Data reflect mean ± SD of n = 18 and n = 14 for stress vitiligo (SV) and non-stress vitiligo (NSV), respectively. (G) mRNA abundance of IFN-II (IFN-γ)-related gene expression.

## Results

### 1. Vitiligo patients with certain aggregation of mental factors have a distinct IFN-I-dependent signal expression profile in the blood.

In vitiligo patients, skin melanocytes are partially or completely lost, and no melanin production is synthesized in this area. The exact cause of the destruction of epidermal or follicular melanocytes is complex and remains not yet fully understood, though there have been several theories (autoimmune, biochemical hypotheses, and neural) (19, 20). Neural theory has a wide range of supportive evidence. Abnormalities in both humoral and neurotransmitter (5-HT, SP, CGRP) have been documented (5, 21, 22). In our study, all vitiligo patients were evaluated with Hamiton Depression Scale (HAMD, 24 items) and Hamiton Anxiety Scale (HAMA) by psychiatrists. When HAMA score was more than 7 or HAMD score was more than 8, these patients were considered to have depressive or anxiolytic characteristics. Following hierarchical clustering analysis, all subjects were divided into two major categories, including vitiligo patients with psychiatry “stress vitiligo (SV), n=18” and vitiligo without psychiatry “non stress vitiligo (NSV), n=14” (**Figure 1A** and **table supplement 1**). In other studies, a questionnaire-based study of 1541 adults with vitiligo was to evaluate the impact of psychological stressors in this patient population (4, 23-25). Psychological stressors should be considered as potential disease triggers in vitiligo patients. Then, we chose 9 stress vitiligos and 9 healthy controls to investigate the gene expression profiles in PBMC (**Figure 1B**). The cDNA microarray analysis showed that thousands of genes were differentially expressed (**Figure 1B**) and the number of down-regulated genes seemed to be larger than that of up-regulated genes. All differentially expressed genes showed 7 distinct systemic signatures (**Figure 1C**). The number of regulated genes in immune system, endocrine system and nervous system was larger than that in other systems (**Figure 1C**), suggesting the activation of the above three systems in stress vitiligo patients. These results also indicate that vitiligo is rather a systemic than a local skin disease, which is involved in immune, neural and biochemical mechanisms. It is also well-known that auto-immunity response has been strongly implicated (20). Higher frequencies of circulating antoantibodies have been observed in patients with vitiligo (26). Therefore, in these patients, anti-melanocyte antibodies are detected. As shown in Figure1-figure supplement 1A-B and **Table supplement 3**, there is no difference of anti-melanocyte antibodies between SV and NSV, suggesting that SV is also related to antoimmunity response. GO analysis and the hierarchical cluster analysis demonstrated that the circulation of stress vitiligo displayed the activation of IFN-I-dependent genes (**Figures 1D-E**) and the increase of serum IFNα (**Figure 1F**). It did, however, slightly affect the expression of IFN-II-dependent genes (**Figure 1G**). Together, the data indicate that psychiatric disorder is considered to be a key cause of vitiligo, which is closely related to the IFN-I-dependent response.

### 2. Chronic stress mice model displayed vitiligo-like phenotype which is closely associated with the indirect IFNα effect.

We as well as other researchers presented evidence that psychiatric stress is an important factor responsible for vitiligo in mice and in human (5, 21, 23, 27). Therefore, we established two types of chronic stress (CRS, CUMS)-induced depressant phenotype and then examined whether chronic stress in animal could affect melanogenic function. Stressed mice displayed depressive-like bahaviors in force swim test (FST) (data not shown). Also, serum corticosterone (CORT) was significantly increased and serum 5-HT was decreased in response to stress (data not shown). Meanwhile, stressed mice presented a significant decrease of weight gain (**Figure 2-figure supplement 2A**). Depressive mice were accompanied by obvious cutaneous whitening (**Figure 2A**). Hematoxylin-eosin (HE) staining result revealed that more black pigment granules were seen in ctrl mice (**Figure 2B**). To determine whether these differences did correlate with circulating IFNα level, we collected blood from stressed mice based on the presence or absence of vitiligo. The serum IFNα levels tended to increase (**Figure 2C**), whereas the local IFNAR expression was lower (**Figure 2-figure supplement 2B**). In Hydroquinone (HQ)-induced classical vitiligo model, the serum IFNα levels were not affected (**Figure 2-figure supplement 2C**). It led us to the belief that IFNα is a systemic effect on the onset of vitiligo but not a local, especially stress vitiligo. Consequently, we wanted to identify whether local IFN-I signals in the skin could directly affect melanogenesis. As shown in **Figure 2-figure supplement 3**, in both the presence or absence of α-MSH, IFNα failed to influence melanin production. However, IFNγ could inhibit basal and α-MSH-induced melanogenesis in B16 melanoma cells and normal human melanocytes (28) (**Figure 2-figure supplement 4**). These findings suggest that mental stress contributes to the hypopigmentary disorders in C57BL/6 mice, which is related with indirect systemic effect of IFNα but not direct local effect.

**Figure 2.**
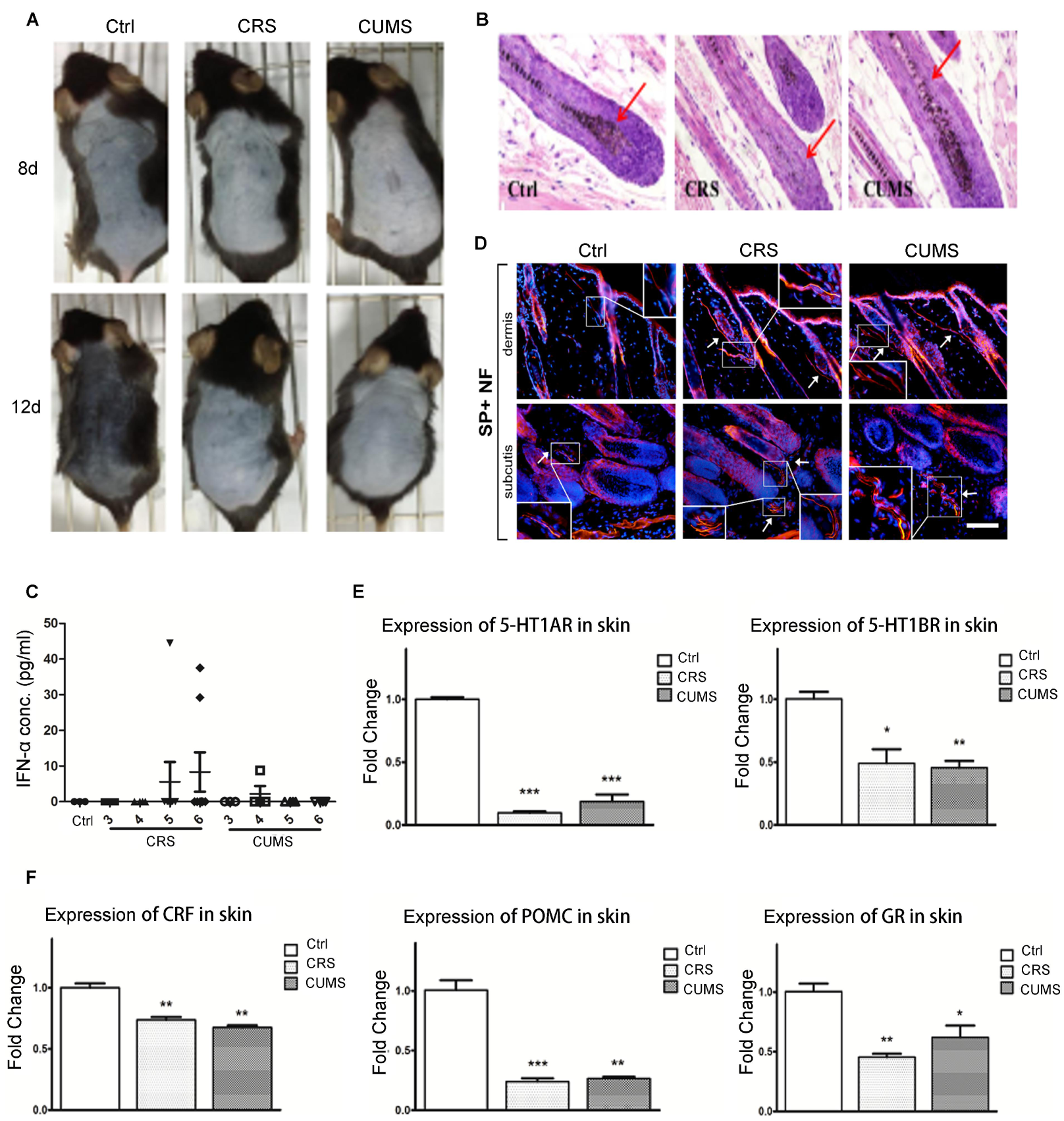
Chronic stress mice model displayed vitiligo-like phenotype which is closely associated with the indirect IFN-α effect. (A) Macroscopic observations of the pigmentary response and the hair cycle stage after stress. The significant area of color in the dorsal skin was from neck to tail. Representative images from 10 animals are shown. (B) A representative area of each group on day 12 after depilation with the majority of hair follicles. Representative images from 3 animals are shown. Original magnification was ×400. (C) Effect of stress (CRS and CUMS) at week 3, 4, 5 and 6 on the serum IFN-α level. Data reflect mean ± SD of n = 3 (D) Effect of stress (CRS and CUMS) on the expression of the cutaneous SP positive nerve fibers. Representative images from 3 animals are shown. Original magnification was ×200. (E) Effect of stress (CRS and CUMS) on the expression of the cutaneous 5-HT1A/1B receptor. Data are presented as mean ± SD, n = 7 in each group, \**P* < 0.05, \*\**P* < 0.01 and \*\*\**P* < 0.001 vs control group with ANOVAs followed by post hoc Turkey test. (F) Effect of stress (CRS and CUMS) on the expression of the cutaneous HPA-axis elements (CRF, POMC, and GR). Data are presented as mean ± SD, n = 7 in each group, \**P* < 0.05, \*\**P* < 0.01 and \*\*\**P* < 0.001 versus control group with ANOVAs followed by post hoc Turkey test.

Previously, we have also found that cutaneous local 5-HT-5-HT1A/1B system, SP/NK1R system and HPA axis were involved in stress-induced depigmentary response (5, 21, 29). We further detected 5-HTR1A/1B, SP and HPA axis expression in skin. As expected, stress administration resulted in the increase of SP-positive fibers and the decreased mRNA expression of 5-HT1A/1B receptor and HPA-related elements (corticotropin-releasing hormone, CRF; pro-opiomelanocortin, POMC and glucocorticoid receptors, GR) (**Figures 2E-F**).

### 3. IFN-α (i.c.v) induced stress vitiligo symptoms through receptors IFNAR and 5-HT1AR

Only a portion of depressive mice developed vitiligo and these mice showed a tendency to higher IFNα in blood. Shiozawa et al (30) investigated in more detail the relationship of cerebrospinal fluid (CSF) and serum IFNα to lupus psychosis and claimed that IFNα was a mediator of neuropsychiatric syndromes in SLE (30). As mentioned, psychiatric vitiligo is a systemic disease and is related to circulating IFNα levels. We next investigated whether higher frequencies of IFNα in brain or in blood could predict stress vitiligo. i.c.v.-injected IFNα contributed to the co-occurence of depressive-like behavior and vitiligo phenotype. The central nervous system (CNS) inhibitory effects showed the increased MAO and Nos activity and the decreased AchE activity in serum (**Figure 3-figure supplement 5**). IFNα i.c.v. treatment for 7 days reduced the crossing and significantly elevated the immobility time in tail suspension test (TST) and forced swim test (FST) (data not shown). Meanwhile, IFNα-treated mice were characterized by the absence of a black pigment in dorsal coat (**Figure 3A**). Several years ago, we confirmed that IFNα could induce depressive-like behaviors through 5-HT1A receptor (31). *In vivo* i.c.v. administration of neutralizing antibodies to the IFN-I receptor (IFNAR) and of interfering IFN-I signals with 5-HT1A agonist (8-OH-DPAT) to the brain, the above changes (depression and depigmentation) could be restored (**Figures 3A-B** and **Figure 3-figure supplement 5**). To further explore the molecular mechanisms involved in this process, the expression of key regulators of melanogenesis (TYR, TRP1, TRP2 and MITF) was compared by Western Blot analysis. The expression levels of these proteins were markedly decreased in IFNα-treated skin (**Figures 3C-D**). At the same time, 8-OH-DPAT and antibodies neutralizing IFN-alpha/beta R1 ameliorated these inhibitory effects of three melanogenesis regulators TYR, TRP1 and MITF (increasing to 87.81%, 43.66%, 75.70%; 66.05%, 29.55%, 62.29% respectively) (**Figure 3D**).

**Figure 3.**
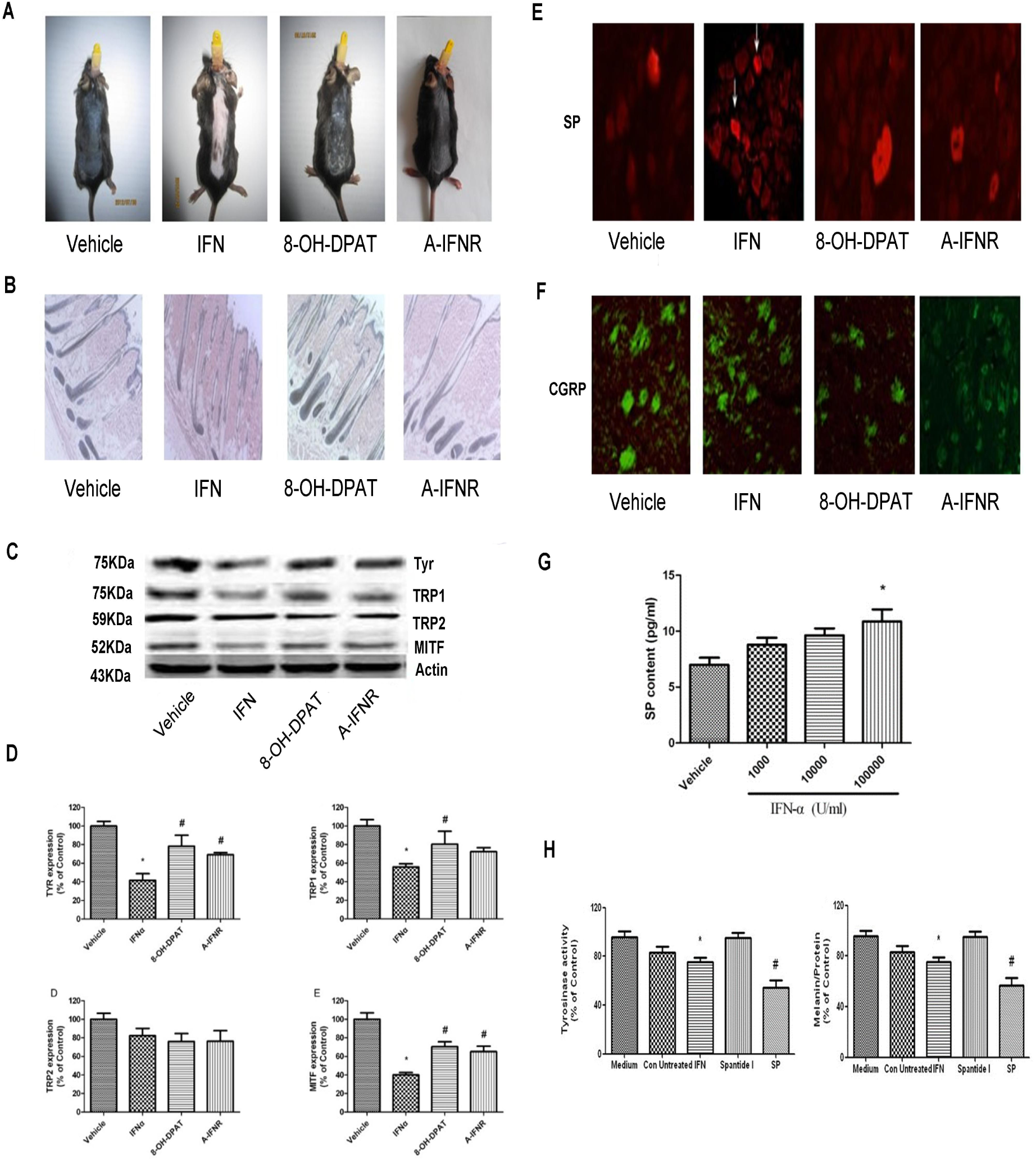
IFN-α (i.c.v.) induced stress vitiligo symptoms through the receptors IFNAR and 5-HT1AR. (A) Effect of IFN-α (i.c.v.) on hair growth and overall hair pigmentation in C57BL/6J mice. Representative images from 10 animals are shown. (B) Effect of IFN-α (i.c.v.) on follicle melanin synthesis in C57BL/6J mice. Representative images from 3 animals are shown. Original magnification was ×100. (C) Effects of IFN-α (i.c.v.) on the expressions of hair follicular TYR, TRP-1, TRP-2 and MITF proteins in C57BL/6 mice. The protein levels of TYR, TRP-1 and TRP-2 were determined by immunohistochemical analysis. Representative Western blots of TYR, TRP-1, TRP-2 and MITF were represented as folds versus vehicle group (D). Expression of β-actin was used as an internal control. Data are expressed as the mean ± SD of individual groups of mice (n=3). Data were analyzed by one-way ANOVA with Tukey’s post hoc test. \**P*<0.05 *vs*. vehicle group, #*P*<0.05 *vs*. IFN-α group. (E) Effects of IFN-α on the expression of DRG SP protein in C57BL/6 mice. SP in DRG was determined by immunohistochemical analysis. Representative photomicrographs of 6 mice skin sections are shown. Cell nuclei are counterstained by DAPI (blue fluorescence). (F) Effects of IFN-α on the expression of DRG CGRP protein in C57BL/6 mice. CGPR in DRG was determined by immunohistochemical analysis. Representative photomicrographs of 6 mice skin sections are shown. Cell nuclei are counterstained by DAPI (blue fluorescence). (G) The effects of IFN-α on the SP release from cultured rat DRG cells. The DRG cells were treated for 24h with various concentrations of IFN-α (100-10000 U/ml) and the SP release was determined by ELISA reagent kit. Data are expressed as means ± SD (n=6). (H) Effect of conditioned medium of IFN α-treated DRG on TYR activity and melanin content in B16F10 cells. Data are expressed as means ± SD (n=6). Data were analyzed by one-way ANOVA with Tukey’s post hoc test. \**p*<0.05 vs Con Untreated group, #*p*<0.05 vs IFN-α group.

Skin is innervated primarily by sensory nerves and by postganglionic parasympathetic and sympathetic nerves (32). Some studies suggest that the nervous system may participate in the maintenance of the physiological integrity and environment of the skin (33, 34). Sensory nerves have been shown to function not only as an afferent system to deal with stimuli from the skin to the central nervous system, but also as efferent system to stimulate target tissue by secreting several kinds of neuropeptides (NPs) (32). Neurotrophic effects of NPs exist in the nervous tissue. Some NPs, including SP and CGRP, are normally made by both sensory nerves in dorsal root ganglia (DRG) (32). Our group has reported that SP and CGRP can directly or indirectly regulate the melanogenesis (29, 35). Based on the above, we have demonstrated that i.c.v injection of IFNα indeed induced the occurrence of vitiligo-like phenotype and depressive signature. Therefore, to investigate whether SP or CGRP are complicated in IFNα-induced interactions between the nervous system and cutaneous melanocytes, SP- or CGRP-positive neurons in DRG were determined. As shown in **Figure 3E**, there was an obvious increase of SP-positive neurons in IFNα group. 8-OH-DPAT or A-IFNR could contribute to the normalization. However, for CGRP-positive neurons, these groups remained unchanged (**Figure 3F**). To determine whether IFNα exert cytotoxic effects on DRG neurons, an MTT assay was done. As shown in **Figure 3-figure supplemental 6**, there was no significant difference between the control and IFNα-treated group. Then, SP release is found to be a drastic increase in response to IFNα (10000 IU/Ml) in DRG neurons (**Figure 3G**). Then this conditioned medium (IFN-CM) could reduce the melanin production and tyrosinase activity in B16F10 cells (**Figure 3H**). This decrease could be further augmented in the presence of SP, whereas was restored by NK1R antagonist (Spantide I) administration (**Figure 3H**).

### 4. IFNα (s.c.) induced stress vitiligo symptoms through the receptor 5-HT1AR

IFNa is a molecule with a molecular weight of approximately 19 kDa and is there fore hardly able to cross the blood–brain barrier (BBB) and brain–cerebrospinal barrier (36, 37). Here, we explored that periphery IFNα functions on vitiligo which has psychiatric comorbidity using mouse model. When subcutaneously treated with IFNα for 7 days, the sucrose preference (6 MIU/kg) was significantly decreased (31). Meanwhile, at this dose, the immobility time in tail suspension test (p<0.01) and in forced swimming test was longer compared to other doses (0.06, 0.6 MIU/kg) (31). We then used this group (6 MIU/kg) to detect global overview of differentially expressed genes in mouse brain. As expected, the hierarchical cluster analysis demonstrated that the expression of depression-related genes were significantly up-regulated in brain following 7- day injection (**Figure 4D**), suggesting their role in psychiatric disorders. A huge number of genes showed the differential expression in nervous system (**Figure 4E**). Also, this group mouse showed behavioral disorders (**Figure 4-figure supplement 7**) and weight reduction (**Figure 4-figure supplement 8A**). Cutaneous IFN-α R1 expression was increased (**Figure 4-figure supplement 8B**). These above data collectively indicate that IFNα (s.c. 6MIU/kg) can indeed induce depression phenotype. Importantly, on day 7 after depilation, IFNα mice displayed obvious whitening of the dorsal skin in a dose-dependent response (**Figure 4A**). In contrast to IFNα mice, vehicle mice showed progressive darkening of the dorsal coat (**Figure 4A**). Also, more follicle black pigment was seen (**Figure 4B**). Several years ago, we confirmed that IFNα subcutaneous treatment could induce depressive-like behaviors through 5-HT1A receptor (31). To preliminarily explore the molecular mechanisms involved in IFNα-induced whitening process, cutaneous expression of 5-HT1A was analyzed. By western blot, the expression level of 5-HT1A receptor was strongly decreased in a dose-dependent manner (**Figure 4C**). Then, these depressive-like behaviors and depigmentary changes could be successfully blocked by the 5-HT1A receptor agonist 8-OH-DPAT (0.5 mg/kg, i.p., 30 min before IFNα administration) (**Figure 4F** and **Figure 4-figure supplement 7**), suggesting the important role of 5-HT1AR in the development of IFNα-induced depression and vitiligo phenotype.

**Figure 4.**
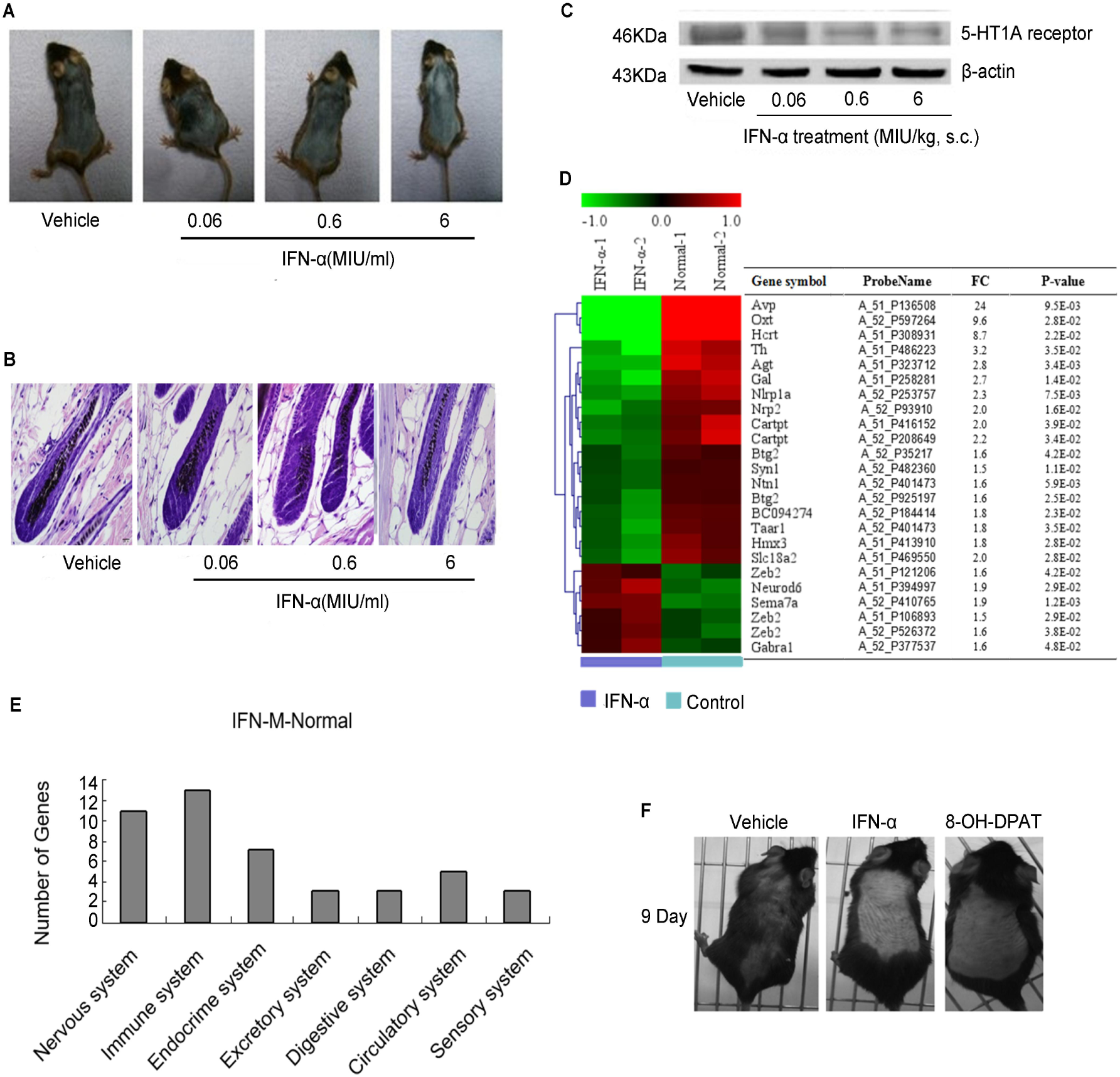
IFN-α (s.c.) induced stress vitiligo symptoms through the receptor 5-HT1AR (A) Effect of IFN-α (s.c.) on hair growth and overall hair pigmentation in C57BL/6J mice in a dose-dependent manner. Representative images from 10 animals are shown. (B) Effect of IFN-α (s.c.) on follicle morphology and melanogenesis of C57BL/6 mice (n=3). Representative photomicrographs of hematoxylin and eosin (H&E). (C) Effects of IFN-α (s.c.) on the expression of 5-HT1A receptor in the skin. Representative Western blot of 5-HT1A receptor from 7 animals was shown. Expression of β-actin was used as an internal control. (D) Heatmap and cluster dendrogram of depreesion-related genes after IFN-α (6MIU/kg, s.c.) treatment. (n=2) (E) Bar graphs of differentially expressed genes in brain nervous system, immune system, endocrine system, excretory system, circulatory system, etc. after IFN-α (6MIU/kg, s.c.) treatment. (F) Effects of 8-OH-DPAT (5-HT1A agonist) on IFN-α-induced depigmentation. Representative images from 10 animals are shown.

### 5. IFNα (s.c.) -induced stress vitiligo symptom is associated with cutaneous 5-HT1A/1B receptor and HPA axis

Experimentally, our group demonstrated that psychology stress (CUMS) could induce the occurrence of the depigmentary process (5, 21). It is mediated by cutaneous 5-HT/5-HT1A/1B system and HPA axis. To ascertain whether peripheral IFNα administration has a similar feature with stressed mice, IFNα were imposed on mice as described in **Figure 4**. The serum corticosterone level was significantly elevated and 5-HT level was decreased (**Figure 5A-B**), suggesting IFNα-treated mice exhibited depression-like symptom. It is well-known that cutaneous 5-HT1A/1B receptor and HPA axis are involved in stress-induced pigment reduction. We further detected skin 5-HT1A/1B receptor and HPA axis-related genes (CRF, POMC and GR) expression following IFNα treatment. Obviously, IFNα administration resulted in the inhibited expression of follicle 5-HT1A/1B receptor (**Figure 5C-D**) and the decreased transcriptional levels of HPA-related elements (CRF, POMC and GR) (**Figure 5E**). To compare with the related depigmentary mechanisms of non-stress vitiligo model (HQ-treated mice), cutaneous 5-HT system and HPA axis are also examined. To our surprise, non-stress vitiligo model mice could not elicit dysregulation of local homeostasis by the skin neuroendocrine system (5-HT system/HPA axis) (**Figure 5-figure supplement 9**). These data suggest that the disturbance of skin 5-HT/5-HT1A/1B and HPA axis induced by IFNα participates in the stress-hypopigmentary processing.

**Figure 5.**
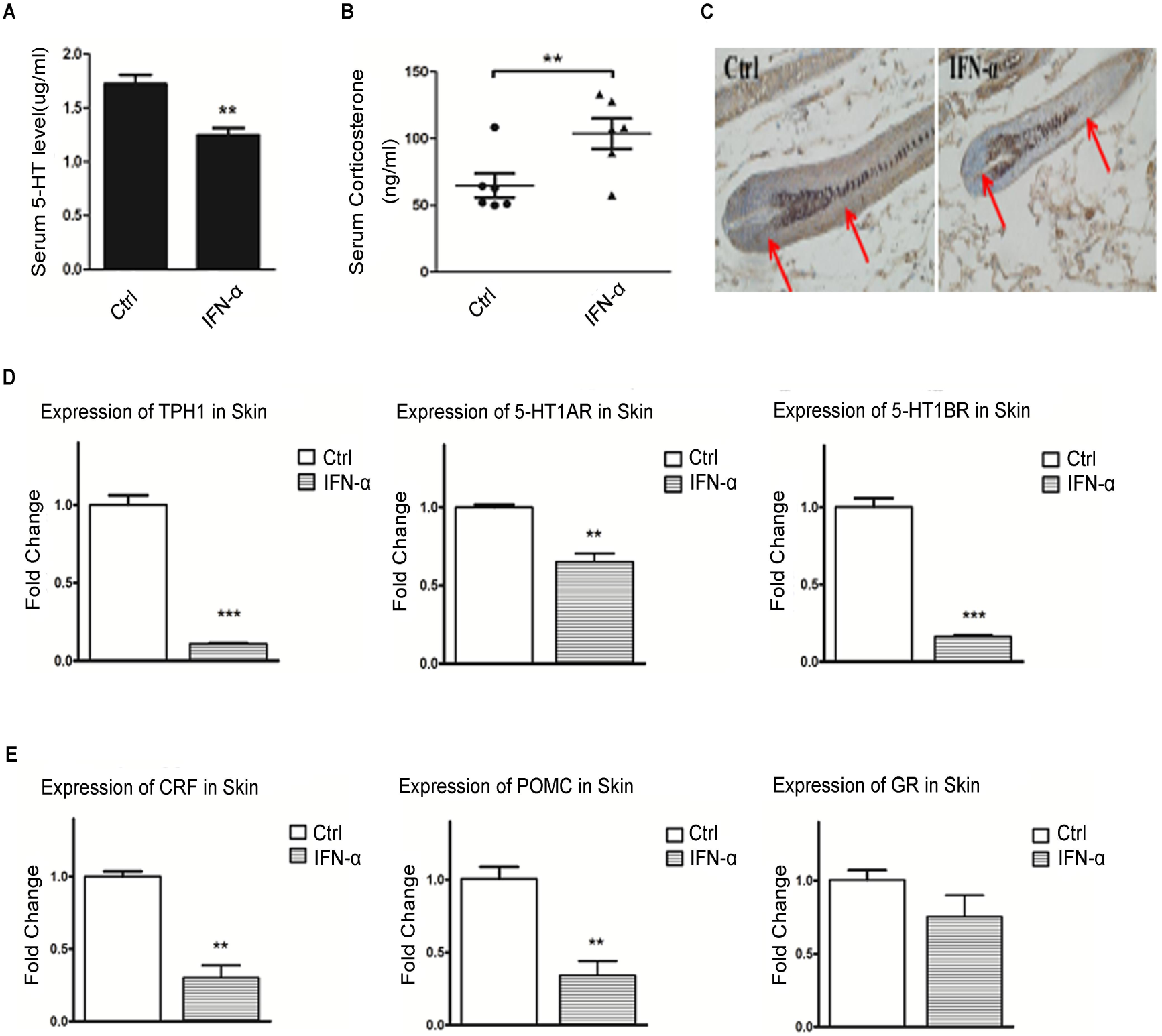
The molecular mechanism of IFN-α (s.c.) -induced stress vitiligo symptoms (A) Effect of IFN-α (s.c.) on the serum 5-HT level. Data are expressed as the mean ± SD of individual groups of mice (n=6). (B) Effect of IFN-α (s.c.) on the serum corticosterone level. Data are expressed as the mean ± SD of individual groups of mice (n=6). (C) Effect of IFN-α (s.c.) on the expression of the cutaneous 5-HT1A receptor. (D) Effect of IFN-α (s.c.) on the mRNA expression of the cutaneous 5-HT-5-HT1A/1B system. (E) Effect of IFN-α (s.c.) on the mRNA expression of the cutaneous HPA-axis elements (CRF, POMC, and GR). Data are presented as mean ± SD, n = 10 in each group. Data were analyzed by one-way ANOVA with Tukey’s post hoc test. \**P* < 0.05, \*\**P* < 0.01 and \*\*\**P* < 0.001 vs control group.

### 6. Type I IFN (IFNα)-related pathway signals in PBMC could discriminate stress vitiligo from non-stress vitiligo.

Vitiligo is often associated with diseases characterized by a type I-IFN signature, such as systemic lupus or psoriasis (38). GO analysis reveals that IFN-I-dependent response and IFNα-related signaling pathway is significantly functional annotated, together with other biological processes including immune system, defense response and innate immune response, etc (**Figure 1D-E** and **Table supplement 2**). These genes also showed a correlation between each other, as shown in the gene regulatory network (**Figure 6A**). To verify whether IFNα-mediated signaling could be used for the classification between stress vitiligo (SV) and non-stress vitiligo (NSV), we firstly measured the levels of IFNα-dependent genes (IFNAR1, IRF7, STAT1, 2,-5,OAS1 and 2,-5,OAS3) by real-time quantitative polymerase chain reaction (q-PCR). It was found that the PBMC of stress but not of non-stress vitiligo induced the expression of IFNα-dependent (IRF7, STAT1, 2,-5,OAS1) genes (**Figure 6B-C**). This response was recapitulated when serum IFNα levels from SV could mostly reach a high concentration (400 pg/ml), which is distinct from the NSV (**Figure 1F**). However, level of type-II IFN (IFNγ)-dependent molecule (39) (e.g., intercellular adhesion molecule 1 ICAM 1), 5-HT, DA, and NE, was both increased or decreased in SV and NSV (**Figure 6D** and **Figure 6-figure supplement 10**). These results indicate that the IFN-I expression program (IFNα) but not IFN-II-associated response (IFNγ) in the circulation could participate in stress vitiligo.

**Figure 6.**
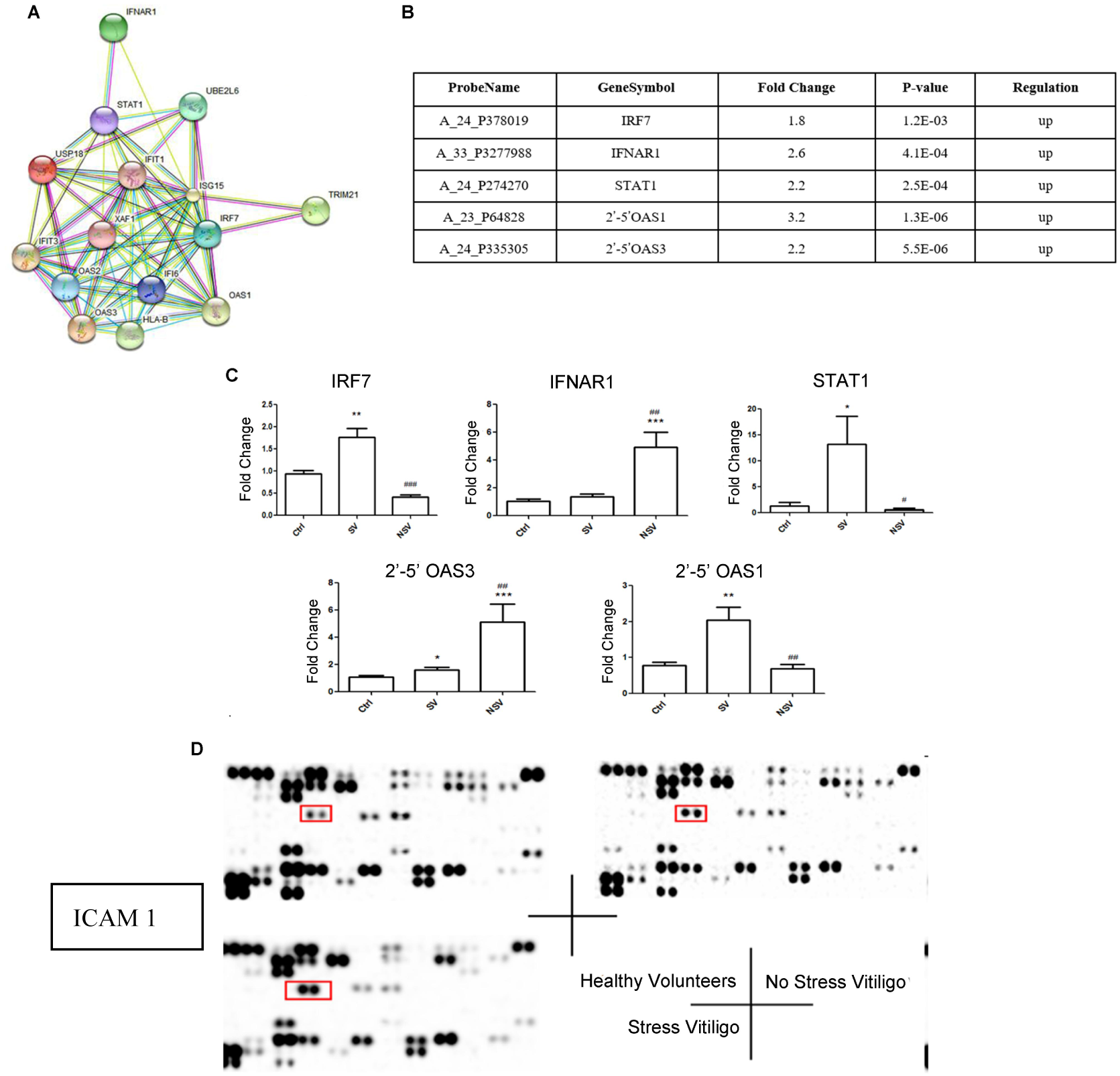
Effects of type I IFN (IFN-α)-related pathway signals in PBMC from the stress vitiligo but not from no-stress vitiligo. (A) Integrated regulatory network of the differential IFN-I dependent gene expression. (B, C) mRNA abundance of IFN-I (IFN-α)-related genes in the PBMC of SV and NSV (fold change relative to ctrl or SV, SV: stress vitiligo; NSV: no stress vitiligo). Data from 32 vitiligo patients and healthy volunteer are expressed in mean ± SD. Data were analyzed by one-way ANOVA with Tukey’s post hoc test. \**P* <0.05, \*\*\**P* <0.001 compared with healthy volunteer group; #*P* <0.05, ###*P* <0.001 compared with stress vitiligo group. (D) Representative images of vitiligo serum immunostained for cytokines and either ICAM 1, type II IFN (IFN-γ)-dependent gene (in box).

## Discussion

Vitiligo is a puzzling disease with a complex pathophysiology. There are three prevailing mechanisms involving the immune, the neural and the autocytotoxic hypothesis (40). In clinical, it is characterized by a silent occurrence and progression in most cases. Before any therapeutic intervention, careful examination under natural light and Wood lamp is needed. However, current examination strategies mainly predict the progressive vitiligo or the stable vitiligo and guide treatment. However, it could not provide accurate therapeutic guide according to disease incentives, thus leading to unsatisfying outcome. Therefore, it is necessary to find a suitable biomarker for vitiligo.

Emerging studies have shown that the progression of vitiligo can be triggered and exacerbated by neural stress (4-7, 25). The neural hypothesis is based on the neurochemistry abnormity in lesion areas, including acetylcholine activity, the distribution of the neuropeptides and the metabolism of catecholamine (41). “Brain-skin axis” concept has aroused more and more attention among researchers. We have demonstrated that mental stresses (CRS, CUMS) indeed induce depigmentation (5, 21) and antidepression drug fluxetine reinforces melanogenesis (42). Therefore, vitiligo can be recognized as a neural systematic disease. In the clinical treatment, vitiligo is only considered to be immune disease. Then, the widely prescribed medication of immunosuppressants eventually leads to low cure rate and extreme side-effects. Therefore, it is urgent to classify the patients according to various pathogeneses, and suit the remedy to the case.

In our research, we aimed at searching for the biomarker to distinguish the psychogenic vitiligo from non-psychogenic vitiligo type. First, according to the gene chip results, the relative gene expression of type I IFN pathway in peripheral blood mononuclear cells of psychogenic vitiligo patients are significantly changed, while the type II IFN pathway are slightly. Also the related gene expression of type I IFN pathway is significantly different between stress and non-stress vitiligo. We established stress-induced vitiligo model. Serum IFNα level was increased in stressed mice. Systematic IFNα could participate in the onset of stress vitiligo through neural, immune and endocrine systems, whereas the local IFNα (skin) could not directly affect the melanin production (**Figure 7**). When the central or peripheral IFNα levels were changed, both of them can simultaneously cause depression and vitiligo phenotype (**Figure 7**). Therefore, IFNα is likely to be a biomarker of psychogenic vitiligo and can be applied to distinguish patients (according to their neural pathogenesis) in the future, and suit the remedy to the case. Mental stress (CRS and CUMS) can trigger depression and depigmentation at the same time, increasing the IFNα level and activating Type I IFN pathway. Moreover, high level of IFNα may be an important trigger of stress vitiligo. Our hypothesis can be proved by the evidence of Type I IFN characteristics in early phase of vitiligo (12) and depigmentary effects of IFNα medication (15). Some researchers suppose that IFNα can be treated as a stressor, and the subsequent manifestations, such as fatigue, inattention and hebetude, are the secondary effects (9, 18).

**Figure 7.**
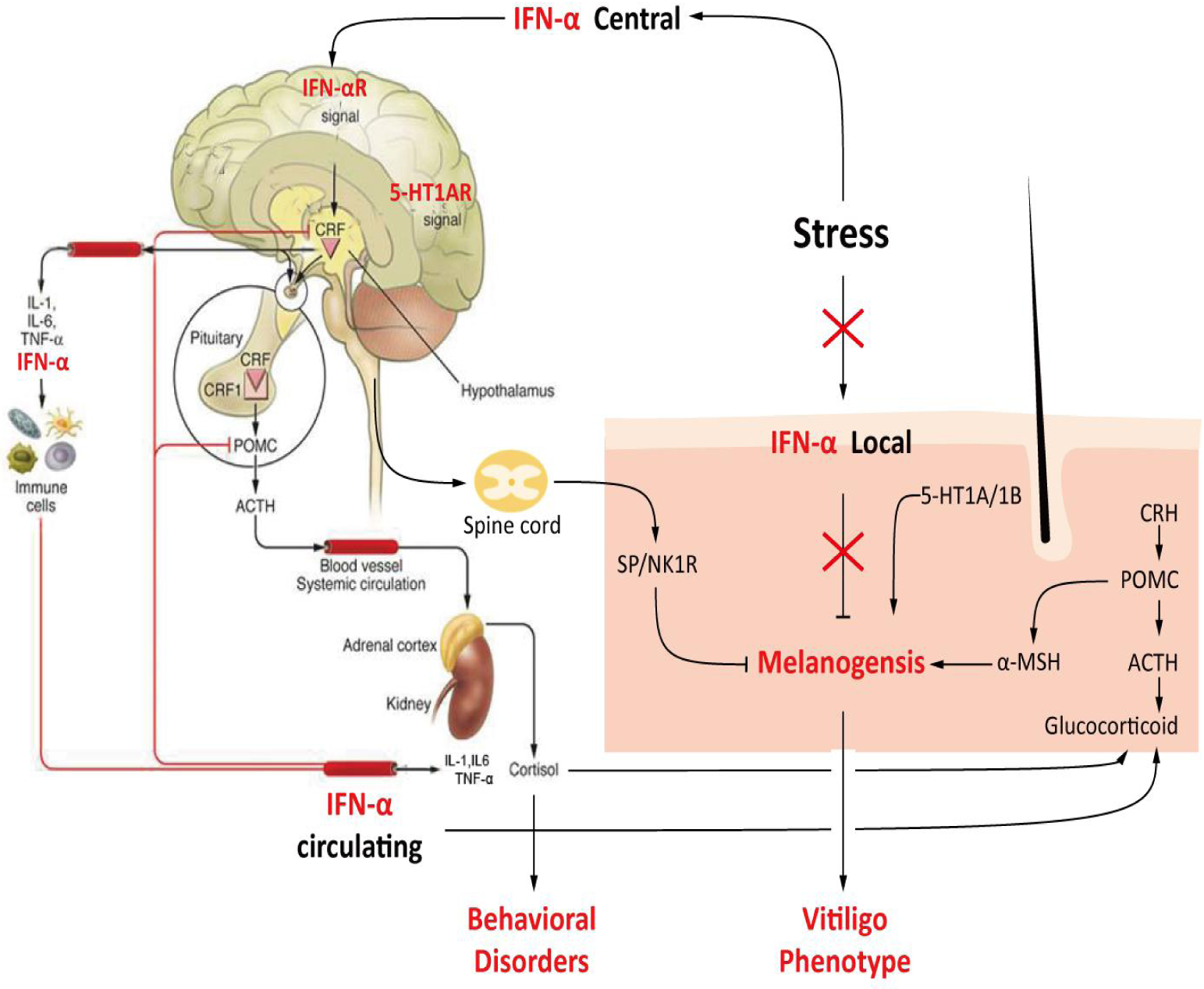

The mechanism of how IFNα regulates melanin synthesis has not been clearly illustrated, whereas the downstream signaling, such as MxA protein and CXCL9, has already been documented (12). IFNα can not directly modulate melanocytes to produce melanin, but IFNγ can influence the maturity of melanosome or directly inhibit the synthesis of melanin granule (11, 12). All the above-mentioned evidences indicate that IFNα may act through an indirect way in the participation of the early pathogenesis of vitiligo, whereas the alteration of IFNγ level may be the subsequent partial effect in lesion areas. The IFNα level in the brain of patients with neurogenic systemic lupus erythematosus is significantly greater than that in blood (30). Only few in periphery are able to cross blood-brain barrier; also, central and peripheral IFNα intervention can cause depression (18). These evidences suggest whether central and peripheral IFNα alteration can lead to depigmetation, or in other words, whether the alteration of IFNα can predict the occurrence of psychogenic vitiligo. Firstly, intracerebroventricular injection of IFNα can lead to the development of depression and vitiligo comorbidity. That pharmacological blockade of IFNAR in brain reversed cutaneous MITF, TYR and TRP1 expression and pigment granules in IFNα administration.

Central IFNα can also affect brain function via secondary effectors such as humoral, NP or cellular components of the peripheral immune system (18). In addition, administration of IFNα can increase the expression of endogenous IFNα in the hippocampus (18). Therefore, both endogenous and exogenous IFNα is able to participate in IFNAR and 5-HT1A signaling pathway, thus resulting in vitiligo-like phenotype. Some researchers found that IFNα plays a key role in stress coping and environment adaptation (18). Under stress conditions or in a competitive environment, IFNAR signaling may be required to maintain Treg homeostasis and function (16). SP and CGRP, as stress factors, are capable of precisely regulating the interaction between skin and nervous system (33). Thus we hope to study whether SP and CGRP are involved in the depigmentation process caused by IFNα. It is well known that the perikarya of cutaneous sensory fibers are localized either in the dorsal root ganglia (DRG) or, those innervating the face and upper neck, in the trigeminal ganglion (33). Ortho/antidromic activation of afferent nerve fibers results in simultaneous signal transduction and release of neurotransmitters (mainly SP and CGRP) at the same site (43). Thus we firstly examined the SP and CGRP-positive neurons at the DRG site after IFNα i.c.v injection. Interestingly, SP-positive neurons were markedly increased in DRG, whereas CGRP-positive neurons were not. Subsequently, *in vitro* studies showed that IFNα stimulated the SP release from DRG cells in a dose-dependent manner. The melanin production and tyrosinase activity were reduced significantly in this conditioned medium (IFNα treated DRG). This effect could be further augmented when adding SP, and be restored by NK1R (SP receptor) antagonist Spantide I.

*In vivo* and *in vitro* data suggest that IFNα i.c.v injection significantly results in simultaneous release of SP at DRG, and promoting the hypopigmentation process indirectly. Furthermore, studies *in vivo* showed that subcutaneous injection of IFNα increased the level in peripheric serum and mediated the simultaneous pathogenesis of depression and vitiligo through 5-HT1A receptor. Here, gene chip technology was used to validate the depression effects induced by IFNα. Depression related genes were up-regulated significantly, including AVP, Sema7a, Zeb2, etc. These genes displayed a distinct distribution in nervous system. Behavioral tests also showed that IFNα markedly increased the immobility time in the tail-suspension test and in the forced-swimming test. Moreover, the sucrose preference was decreased at the same time. These behavioral dysfunctions were restored by 5-HT1AR agonist (8-OH-DPAT). Growing evidence has also shown that IFNα can induce depression (9, 18). In the other hand, nine days after depilation, IFNα-treated mice obviously failed to produce pigmented hair in a dose-dependent manner, while the control group had already recovered. When the dose was up to 6 MIU/ml, all mice exhibited progressive vitiligo-like phenotype. Certainly, through HE assay, macroscopic observations showed that follicular melanin granules were decreased in IFNα-treated skin, resulting in the dorsal whitening. Previously, our group reported that IFNα (system injection) could downregulate hippocampal 5-HT1A receptor expression and induce depressive-like behaviors via 5-HT1A receptor (31). Then Western blot results indicated that IFNα also had inhibitory effects on cutaneous 5-HT1A receptor levels. Moreover, *in vivo*, after receiving 8-OH-DPAT, 5-HT1A receptor agonist, the depigmentary response of IFNα-treated mice was normalized partially. Subcutaneous injection of IFNα appears to be an important mediator of depression and vitiligo, and the precise mechanism by which IFNα exerts should be further elucidated. Our study has found that skin 5-HT system and HPA axis play a vital role in “brain-skin” connection to regulate skin pigmentation (5, 21). Clinically, IFNα therapy was reported to reduce plasma 5-HT levels and activate the HPA axis (9). In our lab, the “brain-skin” connection investigation was always performed on two chronic stress (CUMS, CRS), which provides a very suitable model to study depression and vitiligo comorbidity. Therefore, we detected 5-HT/5-HT1A signaling and HPA axis in mice skin to reveal the underlying mechanism. After systemic injection of IFNα, we discovered that IFNα exerted negative effects on melanogenesis mainly via upregulating corticosterone levels and downregulating CNS 5-HT, cutaneous 5-HT/5-HT1A system and HPA axis.

For one thing, the serum IFNα in stress-induced vitiligo was significantly increased. For another, central and peripheric IFNα could cause depression and vitiligo simultaneously. Finally, when antidepressant fluxetine was subjected to stress-induced depigmentation animals, the increased IFNα level was compromised (data not shown). The serum 5-HT, DA and NE levels were not discriminating between stress vitiligo and non stress vitiligo. The circulating type I IFN (IFNα) pathway-related genes expression in stress vitiligo were significantly discriminating from non-stress vitiligo, while that of type II IFN pathway was not. These evidences indicate that IFNα play a key role in stress vitiligo pathogenesis and further support the great potential of utilizing IFNα as an important biomarker for its diagnosis, which could benefit the subsequent treatment. However, because of clinical sample size limitation, this biomarker needs to be validated in future. We are working with Shanghai No.1 People’s Hospital to try to collect thousands of clinical samples for the verification of the following biomarker.

## Methods

### 1. Experimental design

The study population (n=32) was recruited from Huainan First People’s Hospital of Anhui Province in May 2015. Blood samples for mRNA expression profile microarray analysis were obtained from 9 patients and 9 healthy controls at baseline.

#### Patients and controls

Vitiligo patients had to meet all criteria for inclusion: (1) age (at least 18 years old but less than 65 years old); (2) A subject was excluded if he/she: was taking any other drugs, was pregnant, or had any other skin diseases; (3) All subjects were evaluated with Hamilton Depression scale (HAMD, 24 items) and Hamilton Anxiety Scale (HAMA) by psychiatrists, and were enrolled in the study if Hamilton Anxiety scale score was more than 7 or Hamilton Depression scale score was more than 8. Healthy controls were enrolled among a pool of China Pharmaceutical University student volunteers.

### 2. PBMC preparation and RNA isolation

Blood was collected from 13 volunteers and 19 vitiligo patients between 8:00 and 12:00 in the morning to limit the effect of circadian variation of cytokine production. BD Vacutainer CPT tubes (BD, New York, N.Y., USA) were used to separate PBMCs from other blood cells. The cells were centrifuged at 1,500 g for 30 min at 20°C. After that, blood sera were collected from the top of the PBMCs. Phosphate-buffered saline was used to wash the isolated PBMCs twice, after which they were centrifuged at 190g for 10 min at 20°C. The supernatant was collected and the cells were stored at –80°C until RNA extraction.

Total RNA was extracted using TRIzol reagent (Invitrogen, Carlsbad, CA) and the RNeasy kit (Qiagen, Valencia, CA) according to the manufacturer’s instructions, including a DNase digestion treatment. RNA was measured on NanoDrop-1000 spectrophotometer and quality was monitored with the Agilent 2100 Bioanalyzer (Agilent Technologies, Santa Clara, CA).

### 3. cRNA synthesis and microarray hybridization

Cyanine-3 (Cy3)-labeled cRNA was prepared from 0.5 μg eligible RNA using the One-Color Low RNA Input Linear Amplification PLUS kit (Agilent) according to the manufacturer’s instructions and sent to KangChen Biotech, Shanghai, China for microarray hybridization. Dye incorporation and cRNA yield were checked with the NanoDrop 1000 Spectrophotometer. 1.5 μg of cRNA with incorporation of >10 pmol Cy3 per μg cRNA was hybridized to Agilent Whole Human Genome Oligo Microarrays (G4112A, containing 41,000+ probe sets) according to the manufacturer’s instructions. In total, twelve gene chips were used for 9 patients. The processed slides were scanned and resulting text files extracted from Feature Extraction Software 9.5 (Agilent). A set of 24,315 raw features were taken into sub-sequent analyses after a filtering process for quality and minimum change. Raw data were imported into the Agilent GeneSpring GX software 7.3 and normalized using the Agilent FE one-color scenario (mainly median normalization). Differentially expressed genes were identified through Fold-change screening. Finally, the processed data were deposited in the European Bioinformatic Institute ArrayExpress (http://www.ebi.ac.uk/arrayexpress/) under accession number E-MEXP-2964.

### 4. Bioinformatics analysis

#### 4.1. Differentially expressed probe sets

As mentioned above, multiple groups comparison problems were of interest in our experiment. For identifying significant probe sets, the random-variance model (RVM, commonly used for comparison of more than two groups) F-test [20] was applied to the 24,315 probe sets. Both p-value (< 0.05) and false discovery rate (FDR) <10% were considered statistically significant. A total of 932 microarray probe sets (presenting 478 genes) were identified with this method.

#### 4.2. Hierarchical clustering and series tests of cluster (STC)

To ascertain whether differentially expressed genes among groups were selected correctly, unsupervised hierarchical cluster analysis was done using 478 identified genes. Spearman correlation was used as a similarity measure between samples. Then, gene expression profiles were analyzed using a method called “Series tests of cluster” (STC), which extracts significant patterns by calculating the scores

#### 4.3. Gene ontology (GO) category and pathway analyses

Significant genes in each unique pattern were subjected to GO term using gene ontology project (http://www.geneontology.org/). GO analysis was applied in order to organize genes into hierarchical categories and uncover the co-expression network on the basis of biological process and molecular function. The co-expression network of gene interaction, representing the critical mRNAs and their targets, was established according to the mRNA degree (44). Meanwhile, the significant genes in unique patterns were subjected to KEGG database (http://www.genome.jp/kegg/) and performed on the basis of scoring. In detail, a two-sided Fisher’s exact test and chisquare test were used to classify the enrichment (Re) of both GO and pathway category. The enrichment (Re) was afforded by

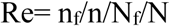

wherein: nf and n represent the number of target genes and total genes, respectively, in the particular GO or pathway, and Nf and N represent the number of genes among the entire differential corresponding target genes and the total number of genes on them GO or pathway, respectively.

### 5. Animals

Adult male C57BL/6 mice (8~10 weeks old, weighing 25-30g) were obtained from the Laboratory Animal Service Center of Yangzhou University. All animals were acclimated for one week under the following conditions: the room temperature was 23 ± 1 °C; humidity was 50 ± 5% with a 12-hour light/dark cycle (lights on at 6:00 a.m. and off at 6:00 p.m.). During this period, food and water were provided *ad libitum*.

#### 5.1 Simple Technique for i.c.v. Injection of Drug.

A cannula for i.c.v. injection of drugs was inserted according to the method of Nakajima et al. (1993) with minor modifications. Mice were anesthetized with chloral hydrate (300 mg/kg i.p.) and placed in a stereotaxic frame (Type 900; David Kopf Instruments, Tujunga, CA). A hole was made through the skull with a needle aimed 0.9 mm lateral to the central suture and 0.4 mm posterior to the bregma. A 24-gauge cannula beveled at one end over a distance of 3.2 mm (Safelet-Cas; Nipro, Osaka, Japan) was implanted into the third cerebral ventricle for i.c.v. injection. The cannula was fixed to the skull with dental cement and capped with silicon. Animals were used experimentally 7 days after implantation.

#### 5.2 Animal Experimental Design and Anagen Induction

Two types of stress, namely chronic restrain stress (CRS) and chronic unpredictable mild stress (CUMS), were imposed on mice. Five mice (Control group) were housed per cage for 21 days. There were 15 mice in every group. According to the reported method, mice (CRS group) were restrained daily for 6 h (10:00 a.m.–16:00 p.m.) before blood and skin samples were collected on day 21 (45). Chronic unpredictable mild stress protocol was adapted from Gamaro et al. (46). On the 9th day of two types of stress, we performed procedures of depilation to induce anagen of hair cycle as described previously (27). In brief, wax/rosin mixture (1:1 on weight) was applied to the dorsal skin (from neck to tail) of C57BL/6 mice with all HFs in telogen. Peeling-off the wax/rosin mixture removed all hair shafts and immediately caused homogeneous anagen development over the entire depilated back area, thus inducing a highly synchronized anagen development.

#### 5.3 Animal Experimental Design (i.c.v. Injection of Drug)

The following drugs were used in the study: IFNα (3SBIO Inc., ShenYang, China), 8-OH-DPAT (Sigma, MO, USA) and Mouse IFN-alpha/beta R1 Antibody (R&D, AF3039, Abingdon, UK). Mice were randomly divided into the following four groups: (1) Control group; (2) IFN-α group: 0.02 MIU/kg of IFNα i.c.v. injection for 7 days (47); (3) 8-OH-DPAT group: IFNα drug injection and application of 8-OH-DPAT i.c.v. injection (6 nmol/0.2 μl) (48, 49); (4) A-IFNR group: application of IFNα concomitant with neutralizing anti-IFNR antibody i.c.v. injection (1.25 μg/μl, 4μl) (50) for 7 days.

#### 5.4 Animal Experimental Design (s.c. Injection of Drug)

The following drugs were used in the study: IFNα (3SBIO Inc., ShenYang, China) and 8-OH-DPAT (Sigma, MO, USA). The first set of mice was subcutaneously (s.c.) injected with IFNα (0.06–6 MIU/kg) for 7 successive days (47), while the second set received 8-OH-DPAT (0.5 mg/kg, i.p.) (51) 30 min before the IFNα administration in a constant volume of 10 ml/kg body weight. Appropriate vehicle-treated (phosphate buffered saline (PBS)-treated) groups were also assessed simultaneously.

#### 5.5 Quantitative real-time polymerase chain reaction (qRT-PCR)

Transcribed cDNA was used as mentioned above. Primers were designed to amplify sequences of 150–250 bp (as shown in **Table supplement 4**). The cDNA samples were used for quantitative real-time PCR analysis. All reactions were carried out on an ABI 7500 Real-Time PCR instrument (Applied Biosystems, Foster City, CA, USA) using the SYBR Green Real-Time PCR Master Mix kit (TAKARA, Japan) according to the manufacturer’s protocol. Amplification conditions were 95 °C for 60s, followed by 40 cycles of 95 °C for 15s and 60 °C for 30s. Each sample was run in triplicate. Glyceraldehyde-3-phosphate dehydrogenase (GAPDH) as the internal control was also amplified under the same conditions to normalize reactions. After completion of the PCR amplification, the relative fold change after stimulation was calculated based on the 2^−ΔΔCT^ method.

### 6. Assessment of Hair Pigmentation

All mice were photographed with a digital camera (Canon, Japan) once every day after depilation. The HE stain was used to quantify the stage of the hair follicles using a published classification technique based on the morphology of the dermal papilla and sebaceous glands (52). In addition, the melanin granule in HFs was visualized histochemically.

### 7. Western Blot

The dorsal skin was quickly dissected out and then lysed in 400 μL RIPA buffer (50 mM Tris-HCl (pH 7.4), 150 mM NaCl, 1 mM PMSF, 1 mM EDTA, 1% Triton X-100, 0.5% sodium deoxycholate, and 0.1% SDS). After centrifugation at 12.000 rpm/min for 20 min at 4 °C, 20 μg of total protein of each sample was loaded into a 12% SDS-PAGE gel and then transferred to PVDF membranes (Millipore). The membrane was blocked with 5% non-fat dry milk in TBS containing 0.05% Tween-20 (TBS-T) for 1 h and incubated with goat polyclonal antibodies against TYR (Product number SC7833), TRP1 (Product number SC10443), rabbit polyclonal antibodies against TRP2 (Product number AB74073, 1:1000, Abcam, Cambridge, UK), mouse polyclonal antibodies against β-actin (Product number CST3700, 1:1000, Cell Signaling Technology Inc., MA, USA). After reaction with the second antibody, proteins were visualized by an enhanced chemiluminescence detection system. Densitometric analysis was again carried out by using the Quantity One (Bio-Rad) to scan the signals. Western blot assay results were representative of at least 3 independent experiments.

### 8. Hematoxylin-eosin (HE) staining

HE staining was performed using an HE staining kit (Solarbio, Beijing, China) according to the manufacturer’s instructions.

### 9. Immunofluorescence

This experiment was performed as previously described with some modifications (53). Sections were dewaxed, rehydrated and immersed in citric acid buffer for antigen retrieval. Then after being washed with 0.01 M PBS, the specimens were treated with PBS containing Tween 20 (PBST) for 15 min at room temperature and then blocked for 1 h in blocking buffer (5% goat serum, 0.1% bovine serum albumin, and 0.1% Triton X-100). Thereafter, the specimens were incubated with each of the primary antibody mixtures (SP, 1: 500, Abcam, ab14184) at 4°C for 24 h. After being washed with 0.01 M PBS, the specimens were incubated with a secondary antibody solution (conjugated goat anti-mouse IgG, Cwbiotech, China) and in the dark inside a cassette at 37 °C for 2 h. The specimens were then washed with 0.01 M PBS and mounted using 50% glycerol and were then observed and photographed under fluorescence.

### 10. ELISA for quantitative detection of human IFNα

Cytokine concentration in serum was measured by commercially available ELISA kits, specific for human IFN-α (eBioscience). Wash microwell strips and Standard dilution on the microwell plate. Then add 80 ul Assay Buffer and 20 ul serum sample and HRP-Conjugate to the microwell plate. Incubate 2 hours at RT. Wash microwell strips and add TMB Substrate Solution Incubate about 10 minutes at RT. Then add stop solution to all wells. The absorbance was measured at 450 nm.

### 11. ELISA for quantitative detection of mouse IFNα

Cytokine concentration in serum was measured by commercially available ELISA kits, specific for mouse IFN-α (eBioscience).Wash microwell strips twice with Wash Buffer.Then add 50 μl of Assay Buffer (1x) to all wells. Add 50 ul of extern diluted standard and Calibrator Diluent and each sample in duplicate to the respective wells. Then add 50 μl diluted Biotin-Conjugate to all wells. Incubate 2 hours at room temperature on a microplate shaker. Wash microwell strips 4 times with wash Buffer. Add 100 μl diluted Streptavidin-HRP to all wells. Incubate 1 hour at room temperature on a microplate shaker. Empty and wash microwell strips 4 times with Wash Buffer. Add 100 μl of TMB Substrate Solution to all wells. Incubate for about 30 minutes at room temperature. Add 100 μl Stop Solution to all wells. The absorbance was measured at 450 nm.

### 12. Proteome profiler Human XL cytokine Array

The Proteome Profiler™ Array (Human XL Cytokine Array Kit) from R&D Systems (Minneapolis, MN, USA) was used to detect the relative levels of cytokines and chemokines for human sera. Briefly, 200 μl of serum was diluted with Array Buffer 6. The sample was added to the membranes, which had already been blocked with Array Buffer 6 and incubated overnight at 4°C on a rocking platform. After three washes (10 min/wash) with 1 x Wash Buffer, the membranes were incubated in diluted Detection Antibody Cocktail and incubated for 1 h on a rocking platform. After three washes (10 min/wash) with 1 x Wash Buffer, the membranes were incubated in diluted streptavidin-HRP for 30 minutes at room temperature. After three washes (10 min/wash) with 1 x Wash Buffer, the membranes incubated Chemi Reagent Mix for 1 minute. Membranes were then exposed to X-ray film for 1 or 10 minutes, and a densitometric analysis of the intensities of the cytokine dots was performed with Image Lab software (Bio-Rad).

### 13. Statistical analysis

All data were expressed as mean ± SD. Statistical analysis of results was performed using one-way ANOVA with Tukey’s correction for multiple comparisons. For comparisons between two independent groups, a Student’s *t*-test was used. *p* < 0.05 was considered statistically significant. Analyses were all per-formed with GraphPad Prism software version 5.0 (GraphPad Software, Inc., La Jolla, CA, USA). All the detailed statistical methods, sample sizes and *p*-values are listed in the **Supplementary statistical information**.

### 14. Study approval

A total of 19 vitiligo patients and 13 healthy controls were enrolled in this research. Written informed consent was obtained from patients and healthy controls before the assessment or measurement began. This research was reviewed and approved by the Ethics Committee of Huainan First People’s Hospital of Anhui Province.

All mice experiments were approved according to the Animal Experimentation Ethics Committee of the Chinese Pharmaceutical University (Approval ID: SCXK-(Jun) 2007-004) and performed in strict accordance with the guidelines of the “Principles of Laboratory Animal Care” (NIH Publication No.80-23, revised in 1996).

## Acknowledgments

The authors would like to thank Chunlei Chen for clinical insights. This work was supported by One Hundred Person Project of The Chinese Academy of Sciences, Applied Basic Research Programs of Qinghai Province (Y229461211); Science and Technology Plan Projects in Xinjiang (2014AB043); Prospective Joint Research Project of Jiangsu Province (BY2016078-02), The National Natural Science Foundation of China (No. 81603216), 2017 CMA-L’OREAL China Skin/Hair Grant (No. S2017140917) and Science and Technology Plan Projects in Qinghai Province (2015-ZJ-733).

## Figure Legends

**Figure 1-figure supplement 1.**
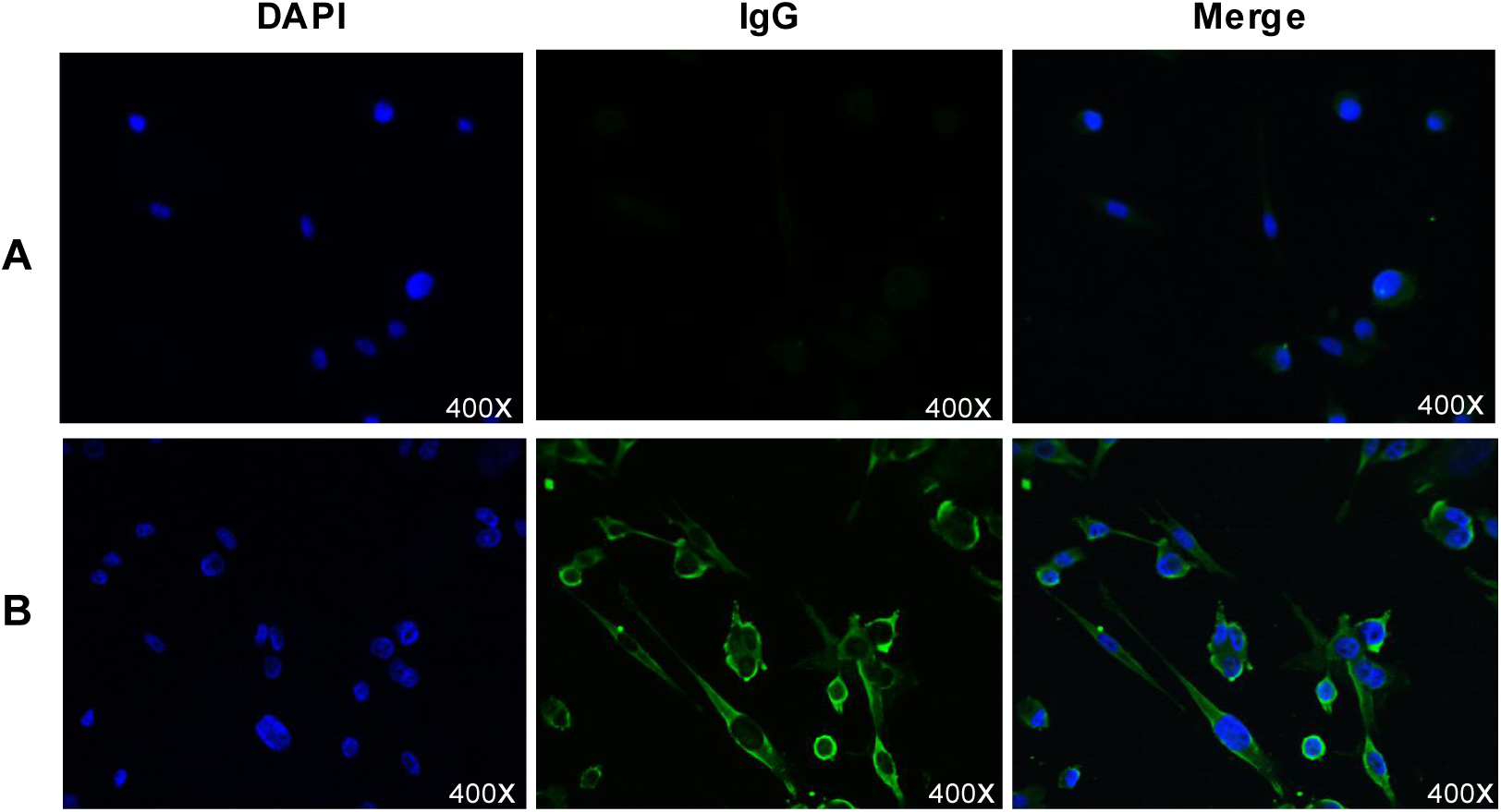
Immunofluorescence detection of serum anti-melanocyte antibodies in patients with vitiligo. A: the test results for the negative cell membrane cytoplasm did not show a green fluorescent expression; B: test results for the positive cell membrane cytoplasm showed significant green fluorescence. Representative images from 32 vitiligo patients and 12 healthy volunteers are shown.

**Figure 2-figure supplement 2.**
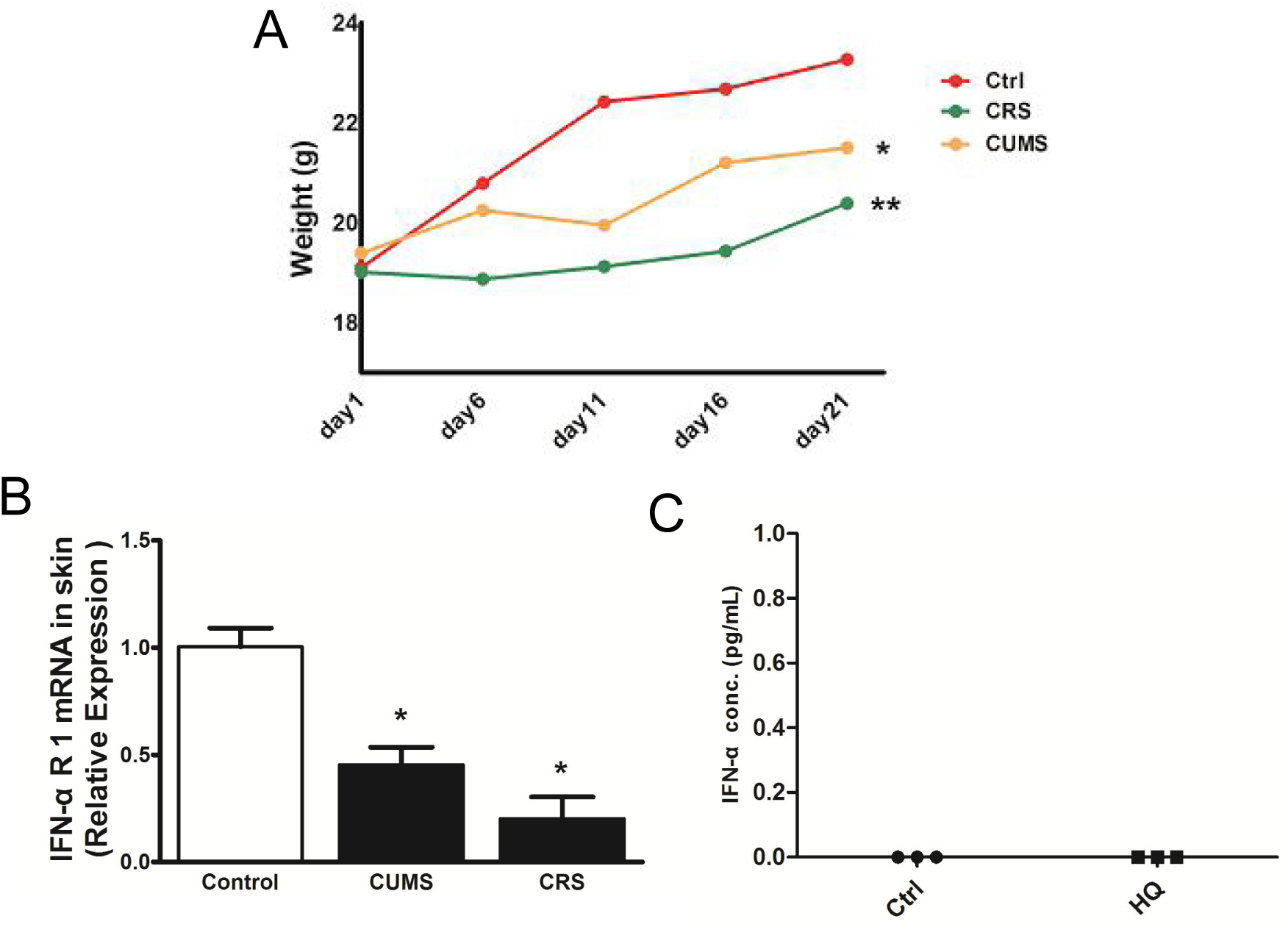
Effect of chronic stress and HQ on cutaneous IFNAR expression and serum IFN-α level. (A, B) Effect of chronic stress on the mice weight and cutaneous IFNAR expression. Data reflect mean ± SD of n = 7. (C) Effect of HQ chemical-induced vitiligo model on the serum IFN-α level. n=3. Data were analyzed by one-way ANOVA with Tukey’s post hoc test.

**Figure 2-figure supplement 3.**
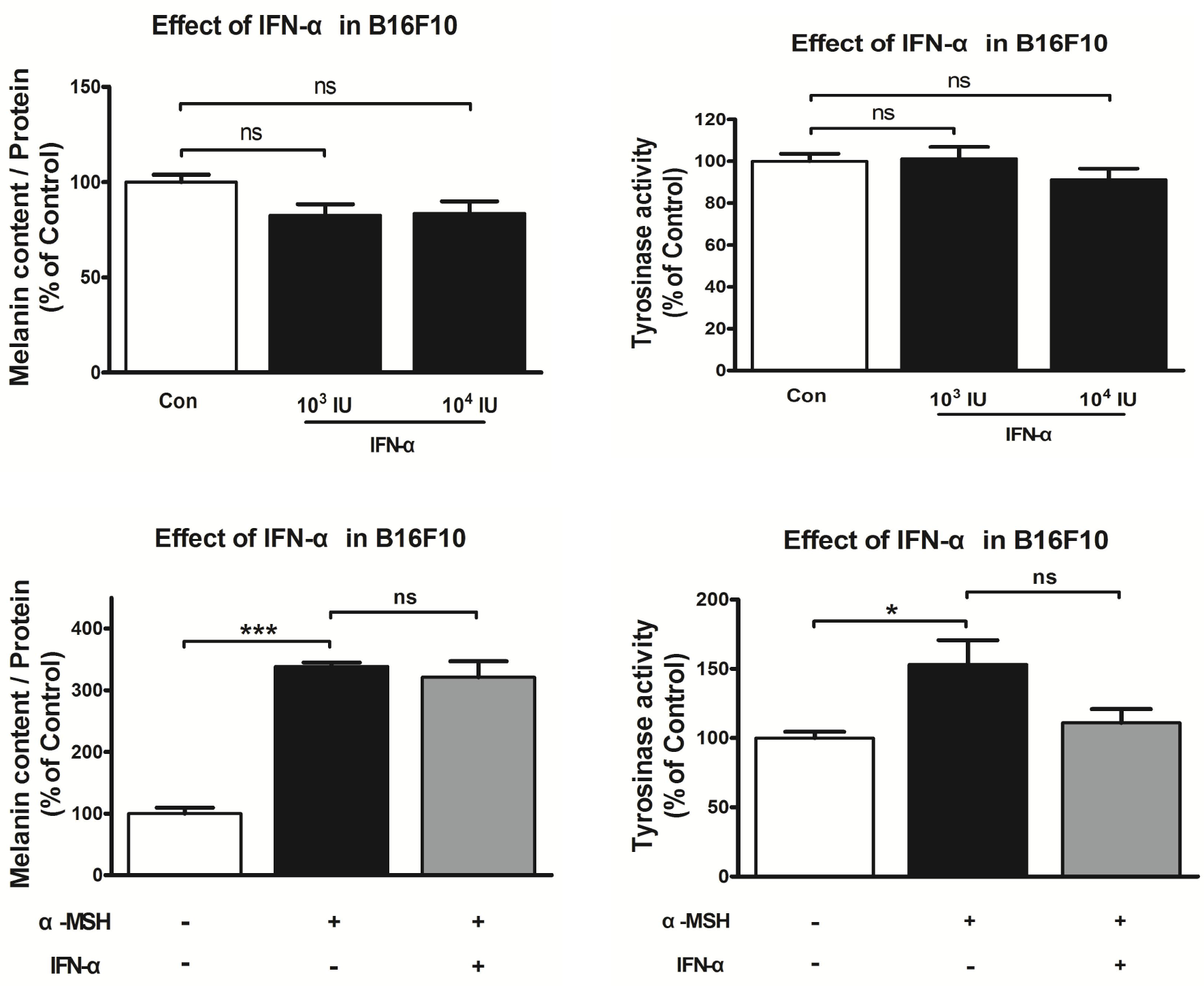
Effect of IFN-α on melanin content and tyrosinase activity in the absence or presence of α-MSH. The B16F10 cells were cultured with IFN-α (1000 or 10000 IU) in the absence or the presence of α-MSH (100 μM) for 48 h. Data are presented as mean ± SD of n=3. Data were analyzed by one-way ANOVA with Tukey’s post hoc test. \**P* <0.05, \*\*\**P* <0.001 compared with the control group; #*P* <0.05, ##*P* <0.01 compared with the α-MSH-treated group

**Figure 2-figure supplement 4.**
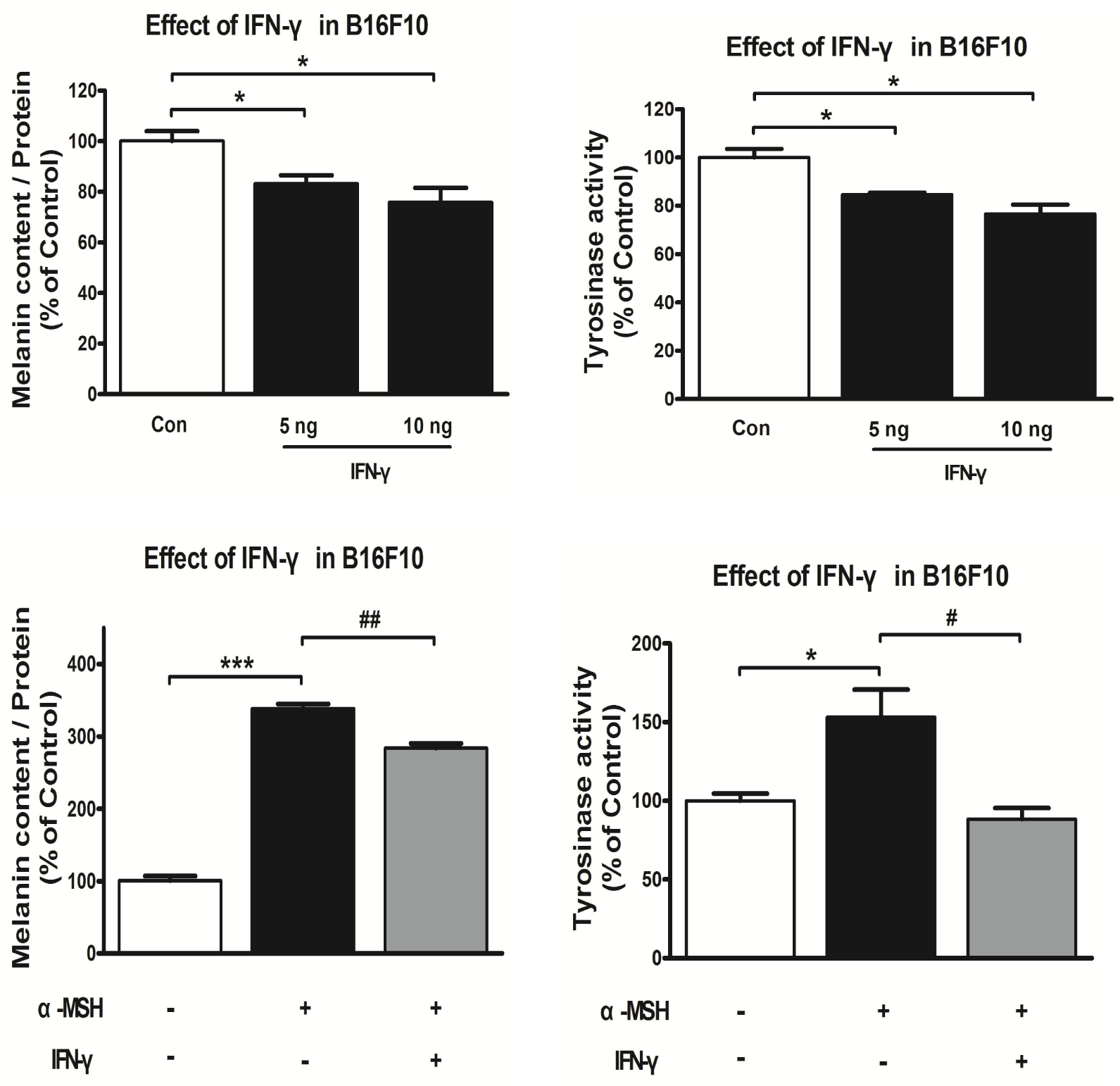
Effect of IFN-γ on melanin content and tyrosinase activity in the absence or presence of α-MSH. The B16F10 cells were cultured with IFN-γ (5 μg or 10 μg) in the absence or the presence of α-MSH (100 μM) for 48 h. Data are presented as mean ± SD of n=3. Data were analyzed by one-way ANOVA with Tukey’s post hoc test. \**P* <0.05, \*\*\**P* <0.001 compared with the control group; #*P* <0.05, ##*P* <0.01 compared with the α-MSH -treated group.

**Figure 3-figure supplement 5.**
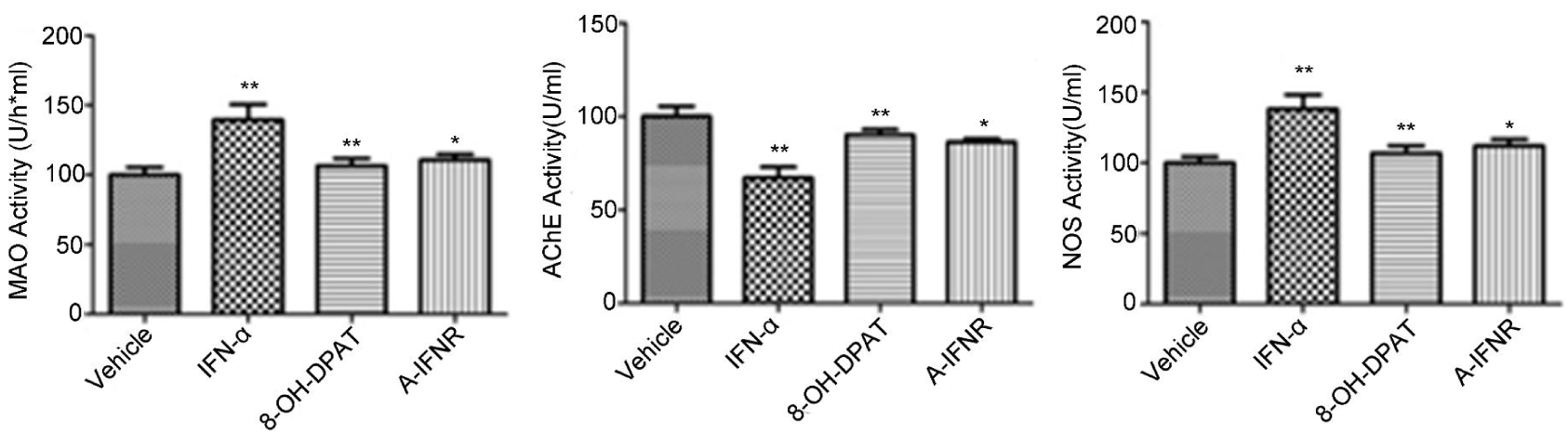
Effect of IFN-α (i.c.v.) on the activity of MAO, AchE and NOS in the serum of mice. Data are expressed as the mean ± SD of individual groups of mice (n=10). \**P*<0.05, \*\**P*< 0.01 *vs*. vehicle group.

**Figure 3-figure supplement 6.**
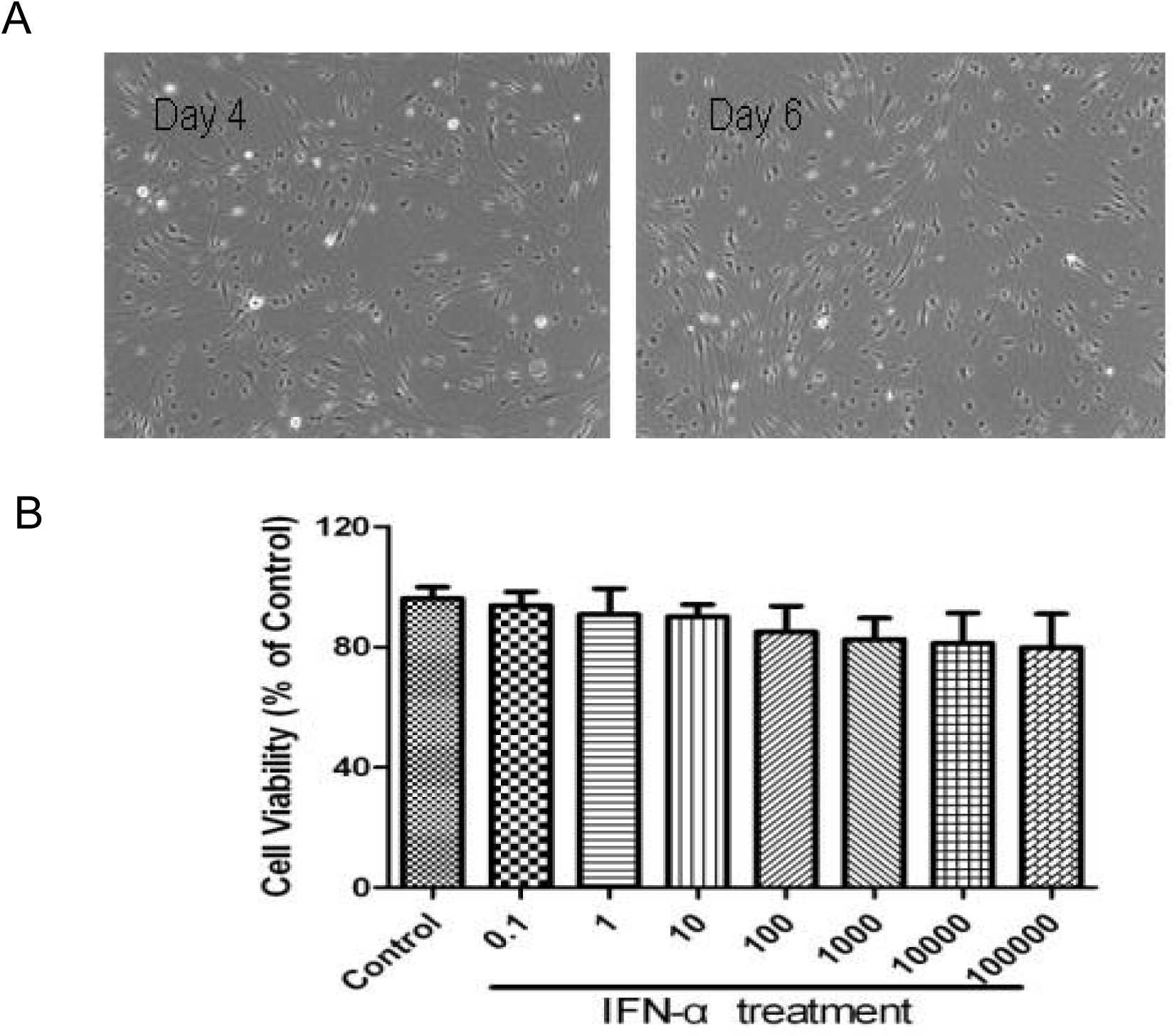
Effect of IFN-α on DRG cell viability. (A) The neuronal culture was generated from DRG of littermates (within 3 days after birth) of SD rats. (B) DRG neurons were treated for 48 h with various concentration of IFN-α (0.1-100000IU/ml) and cell viability was determined by the MTT reduction assay. Data are expressed as means ± SD (n = 6).

**Figure 4-figure supplement 7.**
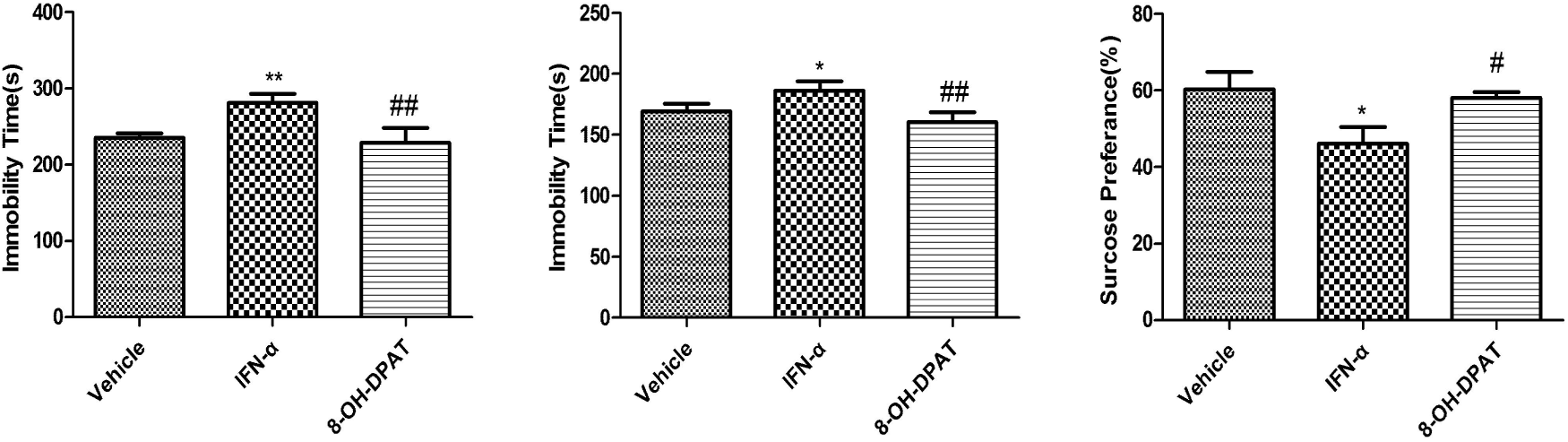
Effect of 8-OH-DPAT (5-HT1A receptor agonist) on the “depressive-like” behavior induced by IFN-α (s.c.) in the forced swim test. (A), the tail suspension test (B) and the sucrose preference test (C). Data are expressed as the mean ± SD of individual groups of mice (n=10). Data were analyzed by one-way ANOVA with Tukey’s post hoc test. \**P*<0.05 *vs*. vehicle group, #*P*<0.05, ##*P*<0.05*vs*. IFN-α group.

**Figure 4-figure supplement 8.**
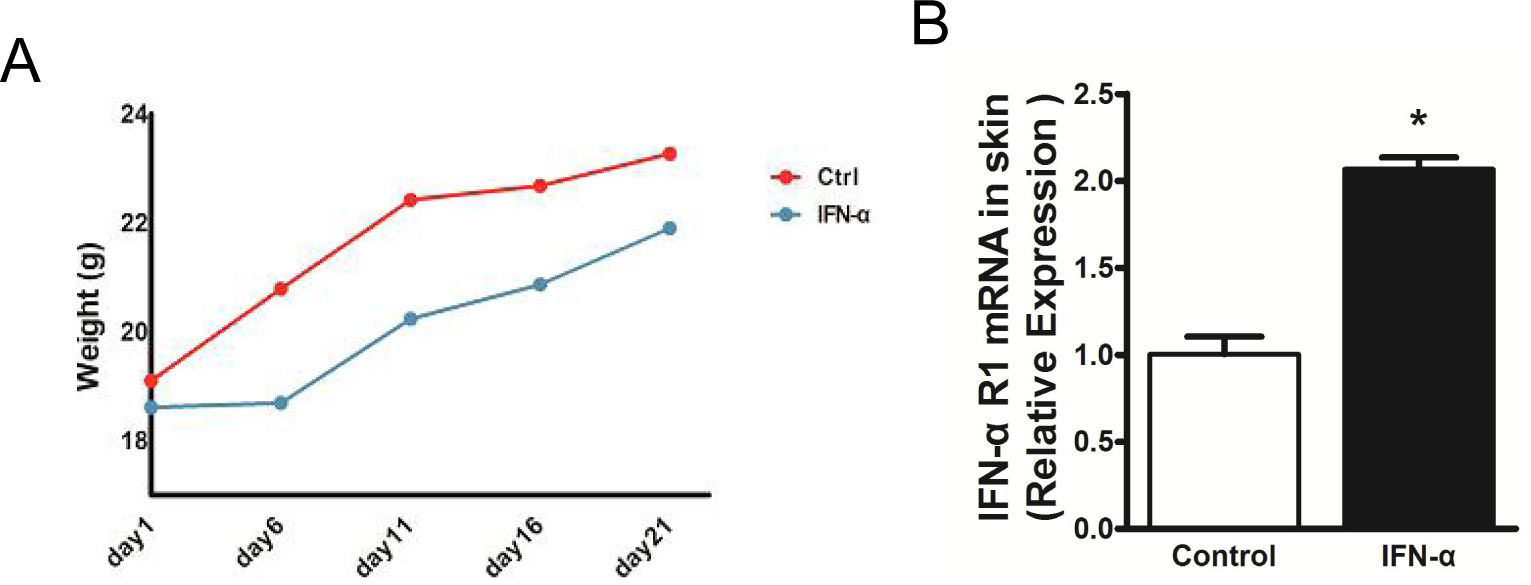
Effect of IFN-α (s.c.) on the weight (A) and cutaneous *IFNAR* mRNA expression (B) in mice. Data are presented as mean ± SD, n = 10 in each group. Data were analyzed by one-way ANOVA with Tukey’s post hoc test. \**P* < 0.05, \*\**P* < 0.01 and \*\*\**P* < 0.001 vs control group.

**Figure 5-figure supplement 9.**
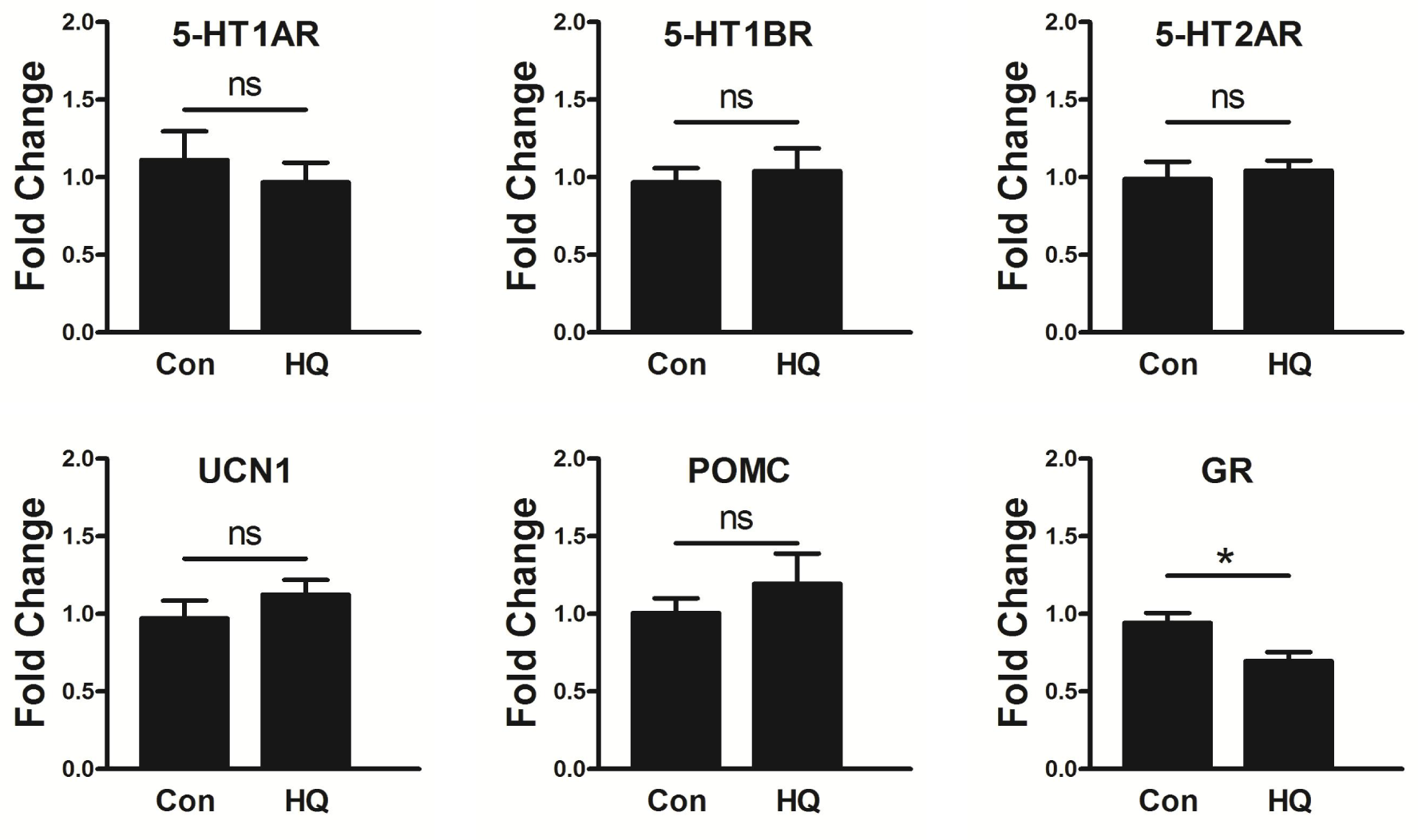
The mRNA expression levels of 5-HT1A/1B/2A receptors and HPA axis elements (POMC, UCN1, and GR) in HQ mouse skin. The expression levels of each gene were normalized against b-Actin then calculated as fold change using the comparative 2^-ΔΔCT^ method. Data are showed in mean ± SD, n = 8. Data were analyzed by one-way ANOVA with Tukey’s post hoc test. \**P*<0.05, \*\**P*<0.01, compared with control (Con).

**Figure 6-figure supplement 10.**
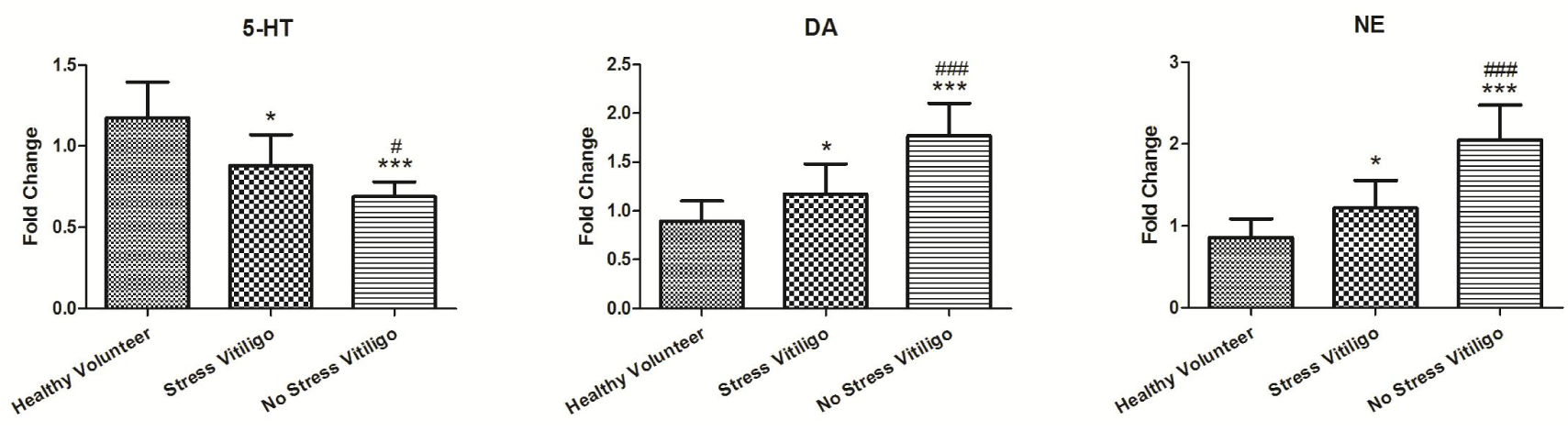
5-HT, DA and NE levels in serum monoamine neurotransmitters in patients with vitiligo. Data from 32 vitiligo patients and 4 healthy volunteer are expressed in mean±SD. Data were analyzed by one-way ANOVA with Tukey’s post hoc test. \**P* <0.05, \*\*\**P* <0.001 compared with healthy volunteer group; #*P* <0.05, ###*P* <0.001 compared with stress vitiligo group.

**Table supplement 1.** Baseline characteristics. Values are mean±SD. HAMA: Hamilton Anxiety Scale. HAMD: Hamilton Depression scale. There are two independently scale. T/t: score. t <7: No anxiety, 14> t ≥7: May have anxiety, 21> t ≥14: Must have anxiety, 29> t ≥21: Must have obvious anxiety,T <8: No depression, 20> T ≥8: May have depression, 35> T ≥20: must have depression.when patients’ Hamilton Anxiety scale score was more than 7 or Hamilton Depression scale score was more than 8 and healthy volunteers’ Hamilton Anxiety scale score was less than 7 or Hamilton Depression scale score was less than 8, will be enrolled in the study.

|  | Vitiligo (n=32) |  | Healthy volunteers(n=16) |  |
| --- | --- | --- | --- | --- |
| Sex | Male(n=15) | Female(n=17) | Male(n=9) | Female(n=7) |
| Age | 35.87±13.84 | 40.35±13.06 | 26.11±2.02 | 25.43±4.79 |
| HAMA Scale score |  |  |  |  |
| t <7 | 6 | 7 | 9 | 7 |
| 14> t ≥7 | 4 | 7 | 0 | 0 |
| 21> t ≥14 | 3 | 1 | 0 | 0 |
| 29> t ≥21 | 2 | 2 | 0 | 0 |
| HDMA Scale score |  |  |  |  |
| T <8 | 9 | 12 | 8 | 5 |
| 20> T ≥8 | 5 | 2 | 1 | 2 |
| 35> T ≥20 | 1 | 3 | 0 | 0 |
| Enrolled | 9 | 10 | 8 | 5 |

**Table supplement 2.**
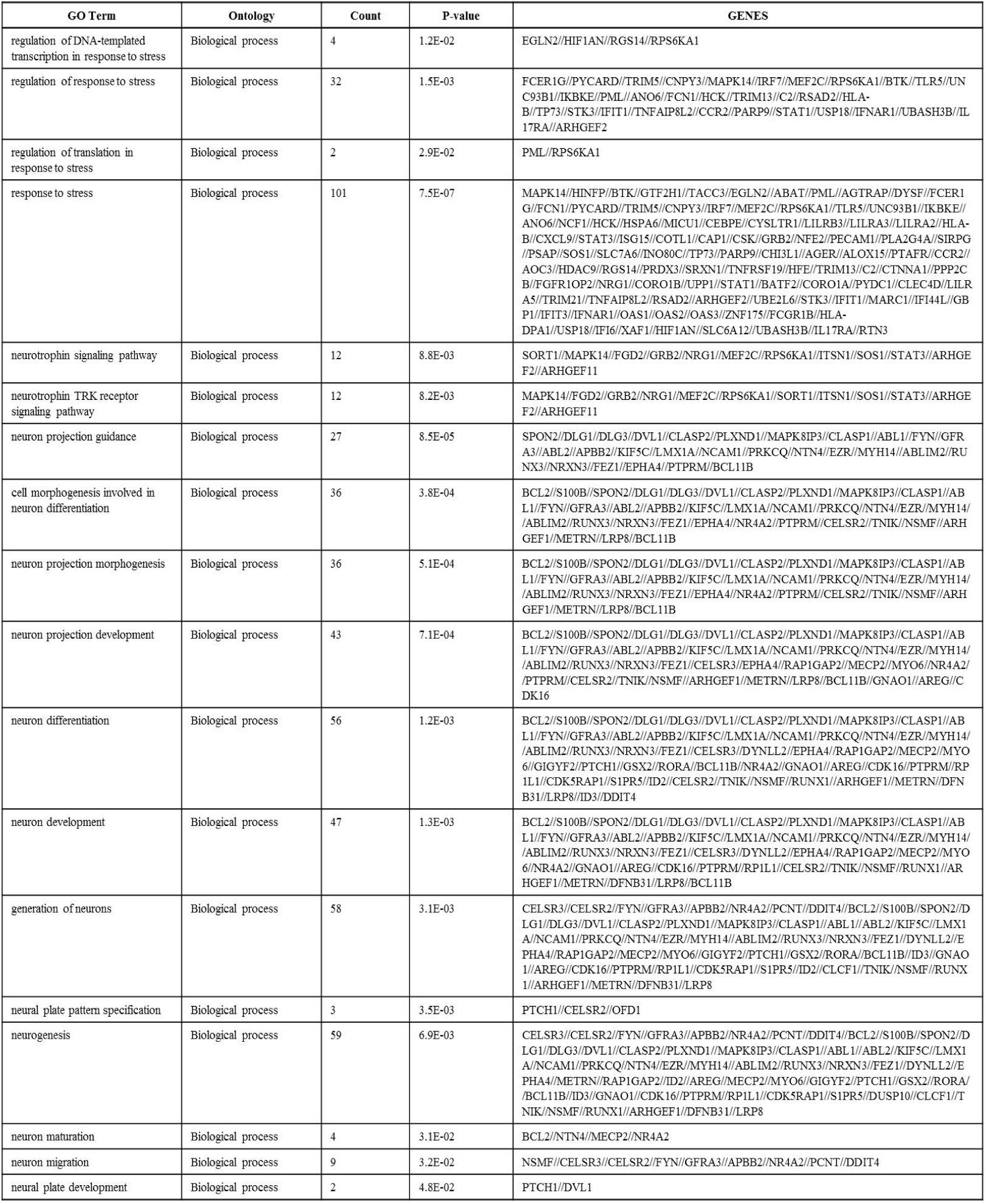
The biological pathways and related genes associated with psychiatric factors were screened by GO analysis

**Table supplement 3.** anti-melanocyte serum IgG antibody positive rate in vitiligo (%)

| classification | cases | 1:10 positive (%) | 1:100 positive (%) |
| --- | --- | --- | --- |
| Stress vitiligo | 19 | 15 (78.95%) | 9 (47.37%) |
| No stress vitiligo | 13 | 13 (100%) | 7 (53.85%) |
| Healthy volunteer | 12 | 4 (33.33%) | 0 (0%) |

**Table supplement 4.**
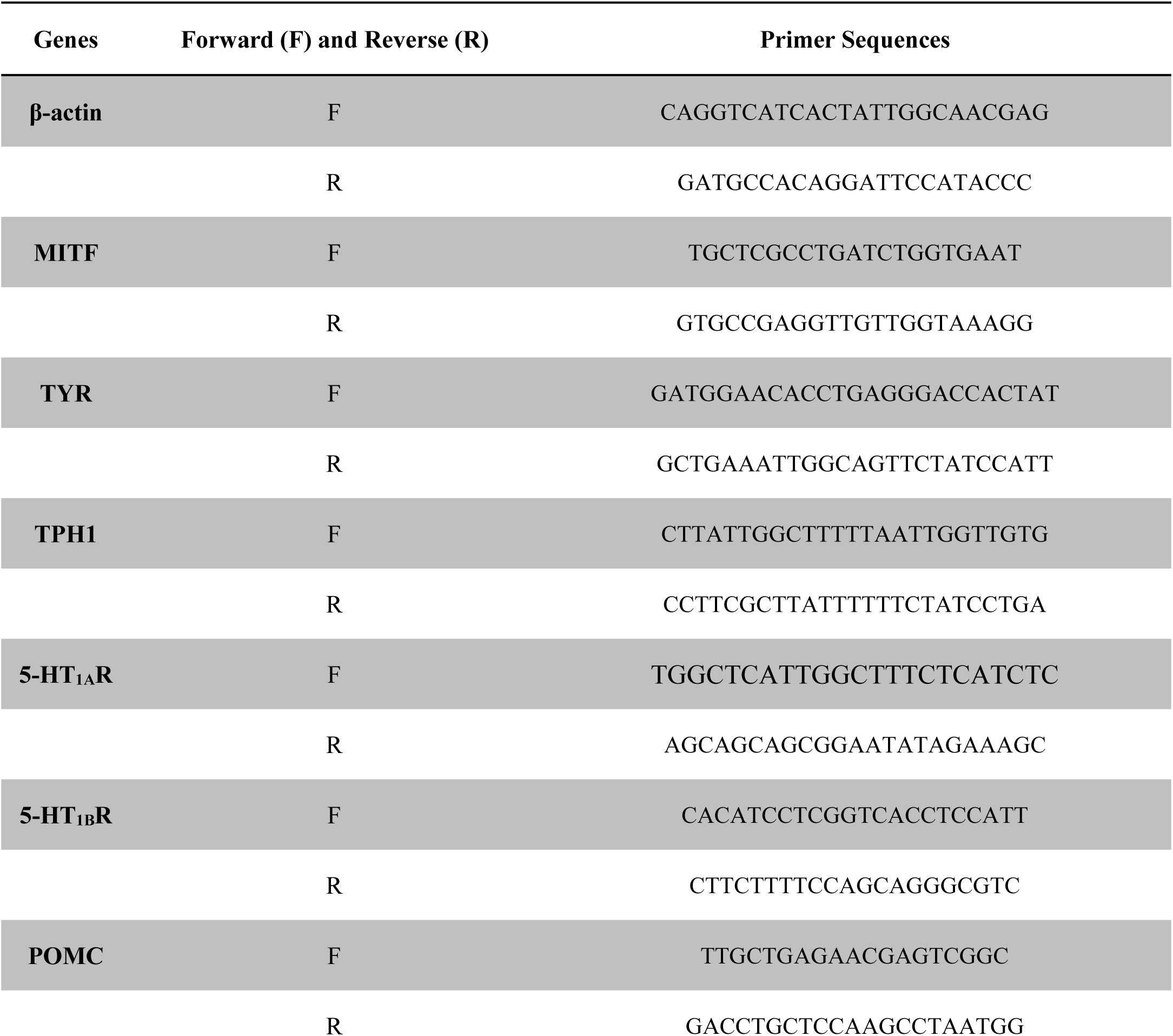
Primer sequences.

## Supplement materials

### 1. LC-MS examination

Chromatographic separations were performed by Finnigan Surveyor LC-TSQ Quantum Ultra AM LC-MS system (Thermo Finnigan, USA) equipped with Xcalibur1.1 workstation, a quaternary pump, an online degasser and a thermostatically controlled column compartment. The mobile phase was composed of (A) water (0.2 % formic acid, 0.1% ammonium acetate, v/v) and (B) acetonitrile. The gradient elution was 2–8% B at 0–6 minutes, 8–70% B at 6–8 minutes, 2% B at 8.01 minutes, 2% B at 8.01–12 minutes. Chromatographic separation was carried out at a Hanbon Lichrospher C18 column (4.6mm × 25cm, 5 μm) with a solvent flow rate of 1.0 ml/minute at a temperature of 30°C. The sample injection volume was set at 10 μl.

### 2. Immunohistochemistry

The fixed brain tissues were embedded in paraffin and were sectioned into 3-μm slices for immunohistochemical staining for the 5-HT1A receptor using an anti-5-HT1A receptor antibody (Abcam, ab85615) (with a maximum of 6 sections per animal). Briefly, sections were pre-treated using heat mediated antigen retrieval with sodium citrate buffer for 20 mins. Then sections were blocked in 5% normal goat serum diluted in phosphate buffer saline (PBS) with 0.25% Tween 20. The rabbit polyclonal antibody against 5-HT1A was used at 1:1000 for 15 mins at room temperature and detected using Horse Radish Peroxidase (HRP) conjugated compact polymer system (54). Diaminobenzidine (DAB) was used as the chromogen. The sections were examined under a laser scanning confocal microscope (Olympus, FV1000).

### 3. Open-field test

The locomotor activity was evaluated as described previously (31). The apparatus was a square, walled arena (50 cm × 50 cm × 22 cm) with white Plexiglas and floor. Each of the mice was placed in the center area of the open-field and analyzed for its motility in a time-period of 5 minutes. During a test period of 5 min, the numbers of crossings (squares crossed with all paws) and rearings (rising on the hind paws) were scored by observers who were blind to the treatment conditions. In this test, the locomotor activity was indicated by the numbers travel in the apparatus while the vertical activity was assigned by number of rearings. At the end of testing, the number of fecal boli was also measured and the arena was cleaned with a 10 % ethanol solution.

### 4. Tail suspension test

The tail suspension test was carried out as previously described (31, 55). Briefly, an adhesive tape was fixed to the mouse tail (distance from the tip of the tail = 2 cm) and hooked to a horizontal ring stand bar placed 30 cm above the floor. All the animals were suspended for 5 min, and the test sessions were video-taped for scoring. Mice were considered immobile only when they hung passively and completely motionless. The immobility time was recorded by observers blind to the treatment conditions.

### 5. Forced-Swim Test (FST)

The forced swim procedure was carried out according to the slightly modified method of Porsolt et al (56). Animals were placed individually into glass cylinders (height 50 cm, diameter 20 cm) containing 30 cm of water maintained at temperature 23–25 °C (57). Animals were allowed to swim for 6 min. After the initial 2 min of vigorous activity, the total duration of immobility was recorded during the last 4 min of the test. Mice were considered immobile when they stopped struggling, remained floating passively, made no attempts to escape and showed only slow limb movements necessary to keep its head above the water. The posture of immobility in the context of the FST was originally coined “behavioral despair” by Porsolt. After each test the cylinder was cleaned. The immobility time was recorded by a trained observer with the help of cumulative stopwatches.

### 6. Measurement of body weight and corticosterone analysis

The body weight of all mice was recorded. Serum corticosterone concentrations were measured using the IBL-AMERICA Corticosterone rat/mouse ELISA kit (IBL, USA) according to the manufacturer’s instruc-tions. Serum samples were incubated at room temperature and then directly used for detection. The lowest detectable concentra-tion of corticosterone that could be distinguished from the “zero calibrator” was 4.1 ng/mL.

### 7. DRG neuron culture

*In vitro* culture of dorsal root ganglion (DRG) neurons was performed according to previously published methods (58, 59). Briefly, adult mice (C57BL/6, 5-6 weeks old, Chinese Academy of Sciences, Shanghai, China) were anaesthetized by I.P. injection of 300 mg/kg chloral hydrate (Sigma-Aldrich, USA) and quickly sacrificed by decapitation. Dorsal root ganglia between L5-L6 spinal segments were extracted and immediately digested by 0.25% Trypsin (Thermo Fisher Scientific, USA) in Dulbecco’s modified Eagle medium (DMEM, Thermo Fisher Scientific, USA) for 30 mins at 37 °C. Trypsinization was stopped with DMEM containing 10% fetal bovine serum (FBS, Sigma-Aldrich, USA). DRG clumps were dissociated by trituration and centrifugation. Neural enrichment was conducted by placing dissociated dorsal root ganglia in a 10 cm cell-culture dish (Corning, USA) containing serum-free DMEM with B-27 supplement (Thermo Fisher Scientific, USA) over night at 37 °C. On second day, floating non-neuronal cell population was removed. The attached DRG neurons were collected and re-seeded in 6-well plate in DMEM containing 10% FBS, B27, Penicillin-Streptomycin-Glutamine (Thermo Fisher Scientific, USA) in a tissue culture chamber with 5% CO2 at 37 °C.

### 8. B16F10 Cell culture

The murine melanoma cell line B16-F10 was purchased from the Cell Bank of the Chinese Academy of Sciences, Shanghai, China and maintained as a monolayer culture in Dulbecco’s Modified Eagle’s Medium (DMEM; Gibco/Invitrogen, Carlsbad, CA) supple mented with 10% (v/v) heat-inactivated fetal bovine serum (FBS; Gibco/Invitrogen), 100 U/ml penicillin, 100 μg/ml streptomycin (Gibco/Invitrogen), at 37 °C in a humidified 5% CO2 incubator.

### 9. Drug treatments

IFN-a (3SBioINS) was dissolved in distilled water before treatment of the samples. To determine IFN-α-induced melanin content, different concentrations of IFN-a (100–10000 U/ml) were evaluated.

### 10. Measurement of melanin content and TYR activity in B16-F10 cells

Total melanin in the cell pellet was dissolved in 1 N NaOH(10% DMSO) for 2 h at 80 °C, and solubilized melanin was measured at 405 nm. Melanin content was calculated from a standard curve using synthetic melanin. Cellular TYR activity was measured according to previously published methods (29, 60).

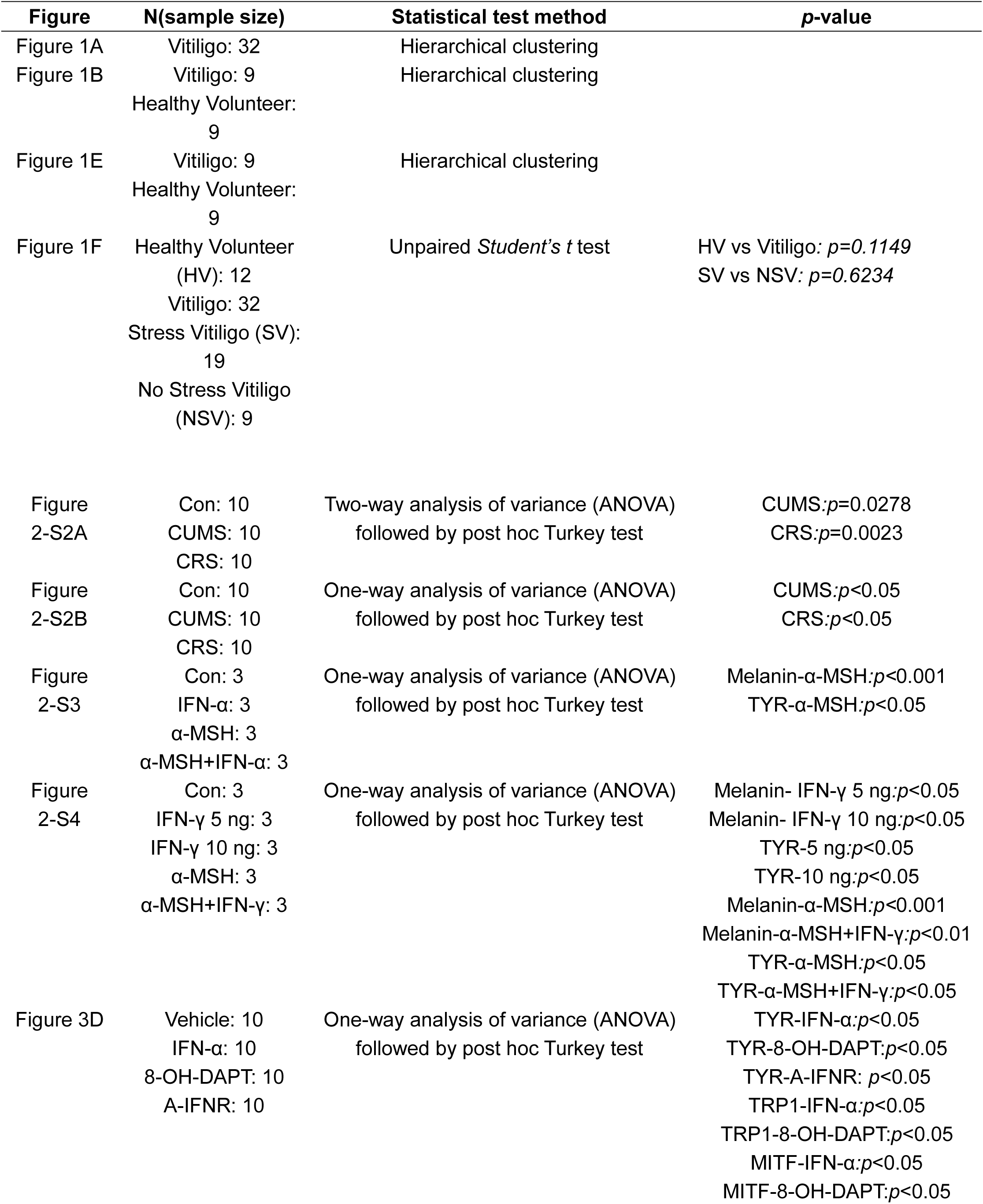
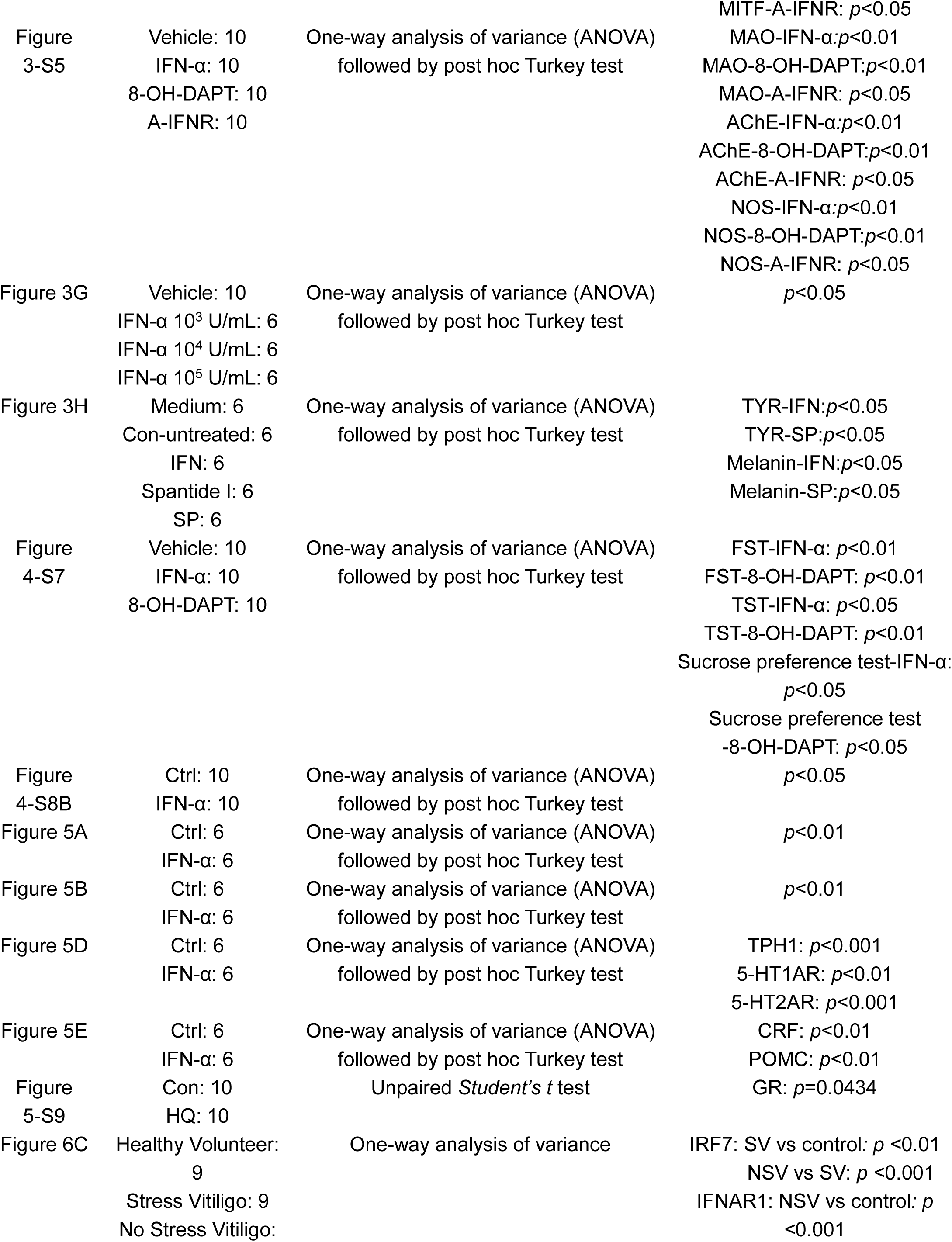
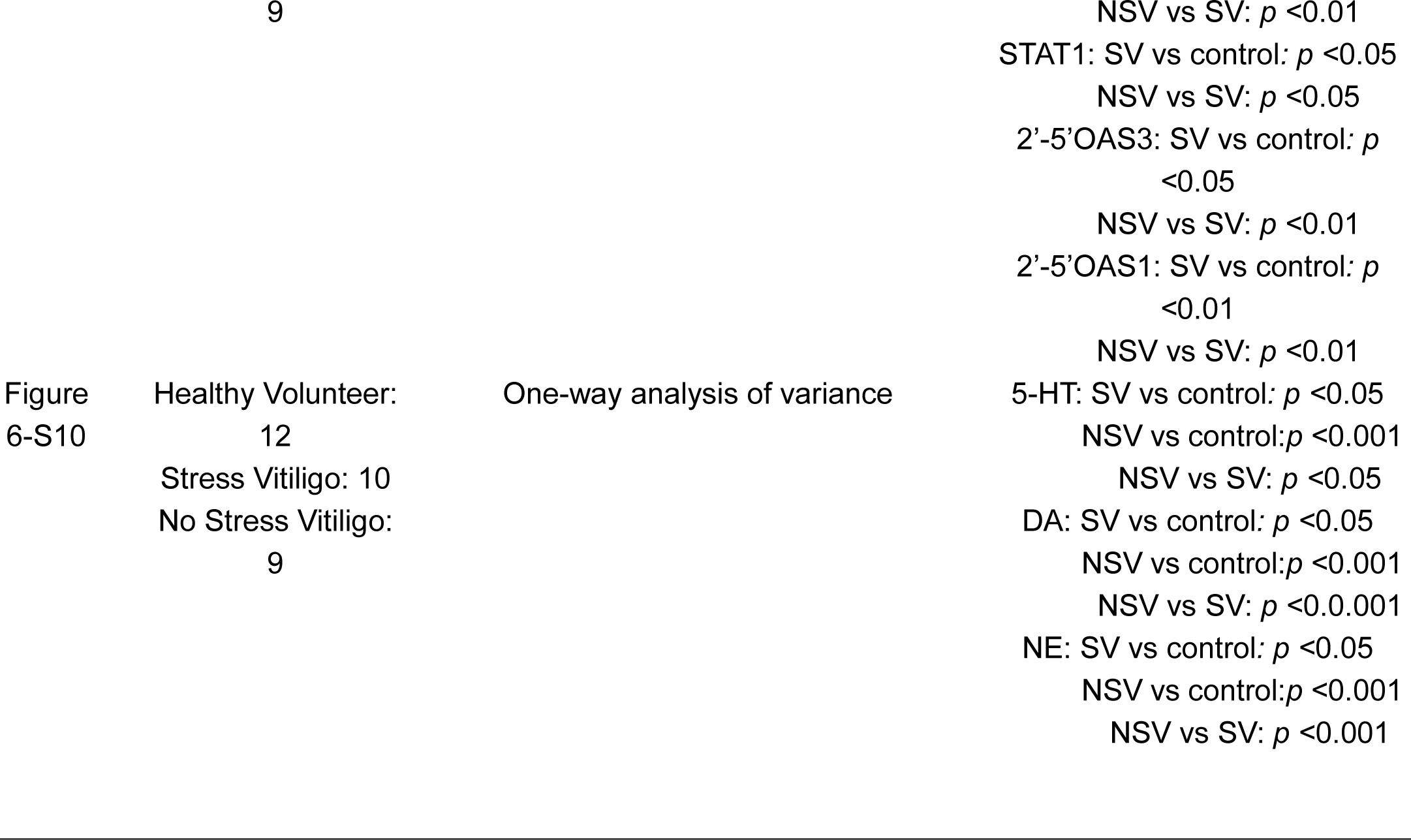
Supplementary statistical information.

